# Uncovering the mechanism of action of a historical antimicrobial against *S. aureus* and *A. baumannii*

**DOI:** 10.64898/2026.02.02.703249

**Authors:** Oluwatosin Qawiyy Orababa, Jessica Furner-Pardoe, Alexander Gale, Blessing Anonye, Joe Ratcliff, Natasha Reddy, Ramón Garcia Maset, Niamh E. Harrington, Shajini Subhaskaran, Séamus Holden, Stephen P. Diggle, Christophe Corre, Freya Harrison

**Affiliations:** School of Life Sciences, Gibbet Hill Campus, University of Warwick, Coventry, CV47AL, United Kingdom; Warwick Medical School, Gibbet Hill Campus, University of Warwick, Coventry, CV47AL, United Kingdom; School of Biological Sciences, Georgia Institute of Technology, North Avenue, Atlanta, GA 30332, United States; Institute of Biomedical Engineering, Department of Engineering Science, University of Oxford, UK (Current address); Institute of Infection, Veterinary and Ecological Sciences, University of Liverpool, L69 3BX, UK (Current address); Centre for Biomolecular Sciences, School of Life Sciences, University of Nottingham, University Park, Nottingham, United Kingdom; Department of Chemistry, University of Warwick, Coventry, CV47AL, United Kingdom; Department of Paleoniotechnology, Leibniz Institute for Natural Product Research and Infection Biology (Hans-Knöll Institute – Leibniz-HKI), Beutenbergstraße 11A, 07745 Jena, Thuringia, Germany (Current address)

**Keywords:** AMR, Ancientbiotics, Antibiotic mechanisms, Antivirulence mechanisms, Biofilm, Natural products

## Abstract

Natural products have provided most of our modern pharmacopoeia, serving as active molecules or scaffolds for active molecules. Their use in drug development is often inspired by their traditional or historical medical use. For many decades, this discovery pipeline has focused on identifying a single molecule responsible for much of the biological activity of a “raw” natural product preparation (e.g. a whole-plant extract) and scoping this molecule for clinical potential. However, it is increasingly realised that historical/traditional remedies with significant biological activity may owe this activity to the combined action of multiple molecules. Concomitantly, microbiologists increasingly argue that effectively fighting antimicrobial-resistant infections will rely on combination therapies that combine multiple antimicrobials and/or adjuvant molecules. We previously reconstructed a complex historical remedy, Bald’s eyesalve. Our reconstruction of this remedy had strong antibiofilm activity, which relied on the presence of multiple ingredients. Here, we report that Bald’s eyesalve has multiple antibacterial effects on exemplar Gram-positive (*Staphylococcus aureus*) and Gram-negative (*Acinetobacter baumannii*) pathogens. Bald’s eyesalve disrupts bacterial membrane integrity; inhibits expression of genes associated with bacterial adhesins, virulence factors and efflux pumps in both *S. aureus* and *A. baumannii*; inhibits quorum sensing in *S. aureus*; and causes downregulation of genes involved in *de novo* nucleotide biosynthesis in *S. aureus*. Lastly, we show that this multifaceted mechanism of action makes it difficult for *S. aureus*, *A. baumannii*, and *Pseudomonas aeruginosa* to evolve resistance against Bald’s eyesalve. Bald’s eyesalve could be used to identify a defined cocktail of natural products suitable for preclinical testing as a multi-target antibacterial preparation to which resistance may arise more slowly than current single-molecule antibiotics.

**Importance:** The increasing mortality and economic cost of antimicrobial resistance (AMR) have made it one of the biggest threats to global health. Available antibiotics are in short supply due to the slow progress in the discovery of new antibiotics. Hence, there is a need for alternative treatment options against difficult-to-treat infections. We have identified a natural product cocktail based on a historical remedy with broad-spectrum antibacterial activity and a multifaceted mechanism of action. We have shown that there is slower resistance evolution to this cocktail compared to mainline antibiotics, and it could be used as the foundation of an alternative treatment to antibiotics in clinical settings.

## INTRODUCTION

In 2021 alone, bacterial antimicrobial resistance (AMR) was associated with 4.71 million deaths (1). If this silent pandemic of increasing AMR persists, close to 170 million lives will be lost to AMR-associated infections between 2025 and 2050 (1). AMR is costly in economic terms as well as in human lives lost: in 2019, total hospital costs and productivity losses associated with AMR infections were close to USD700 bn and USD200 bn, respectively (2). The twin challenges of selective pressure for resistance to mainline antibiotics (3) and the scarcity of new antimicrobial treatments in the clinical (4,5) create an urgent need to find novel therapies with a long clinical shelf life in the face of bacterial abilities to rapidly mutate and adapt.

Increasingly, researchers are hunting for antibacterial agents whose mechanism of action involves attacking multiple cellular targets, and/or that target bacterial structures or functions that are highly evolutionarily conserved, such as the cell membrane. Combination therapies, which comprise two or more active compounds, or which combine active compounds with adjuvants that enhance their activity, are increasingly recognised as one way to achieve this aim (6,7). Combination therapies may range from a simple combination of an antibiotic plus a molecule that reduces the ability of bacteria to evade antibiotic attack (e.g. beta-lactam plus beta-lactamase inhibitor) to a chemically complex preparation which uses several mechanisms to kill bacteria and to which it appears to be hard for bacteria to evolve resistance (e.g. medical-grade honey) (8,9).

Our modern clinical use of honey stems from its long history of use in traditional and “folk” medicine around the world (10,11). Ethnopharmacology and interdisciplinary studies of historical medicine thus have the potential to identify natural products with clinically-significant antibacterial activity. Focussing on plants and plant-derived preparations, it is now clear that many plant extracts contain not just individual compounds with strong antibacterial activity and a promising therapeutic index (12), but combinations of compounds that act additively or synergistically to produce a high level of antibacterial activity (13). A textbook example of a single-molecule antimicrobial compound derived from a plant used in historical medicine is the antimalarial drug artemisinin (14): however, more recent research has shown that complex extracts of the parent plant have more potent antimicrobial activity than purified artemisinin (15).

We and our collaborators in the Ancientbiotics consortium (16) previously explored the antibacterial potential of a historical remedy known as Bald’s eyesalve, from an early medieval English medical text called Bald’s Leechbook (17–19). This multi-part natural product remedy was made by combining garlic (*Allium sativum*), another *Allium* species (we explored onions or leek, *A. cepa* / *A. ampeloprasum*), wine, and bovine bile, and was used to treat a “lump” in the eye, which we interpreted as likely a stye - caused by bacterial infection of an eyelash follicle. Our version of Bald’s eyesalve has been shown to have significant antibacterial efficacy in *in vitro* wound biofilm models and in biofilm-containing biopsies from mice with chronic wound infections. We confirmed that optimal antibiofilm efficacy of this complex natural product remedy was only seen when all ingredients were combined, consistent with synergistic and/or additive interactions existing between the ingredients (17,18). We also showed broad-spectrum activity against several Gram-positive and Gram-negative species (18). Preclinical safety testing revealed no serious adverse effects of the reconstructed Bald’s eyesalve *in vitro* or *in vivo* (19,20).

In this study, we first confirmed that Bald’s eyesalve has good (less than 1/10th of the preparation inhibited bacterial growth) antibacterial efficacy against selected multidrug - resistant and priority pathogens. We then used phenotypic assays and transcriptomics to unravel the mechanism of action of Bald’s eyesalve against *Staphylococcus aureus* and *Acinetobacter baumannii* (WHO priority pathogens for new drug development, with a high AMR-associated mortality burden (1,21). We found that the eyesalve has a multiple antibacterial effects - some of which were unique to one of the two species studied, and some of which were shared. We then used laboratory selection experiments to select *S. aureus*, *A. baumannii*, and *P. aeruginosa* for resistance to the eyesalve, or to commonly-used antibiotics. We found that while all bacteria rapidly evolved high-level resistance to conventional antibiotics, they were unable to evolve resistance to the eyesalve. Bald’s eyesalve could therefore contain an identifiable cocktail of natural products suitable for development into a multi-target antibacterial preparation with clinical potential.

## RESULTS

### Bald’s eyesalve has good antibacterial activity against multidrug-resistant strains of *S. aureus*, *A. baumannii*, and *P. aeruginosa*

We have previously shown that Bald’s eyesalve has potent antimicrobial and antibiofilm activity against a range of bacterial pathogens, and that using onion, rather than leek, as the second *Allium* species gives more consistent activity between batches (18). We prepared fresh batches of eyesalve from garlic, onion, wine, and bovine bile and carried out broth microdilution assays in cation-adjusted Müller-Hinton broth (caMHB) to calculate the minimum inhibitory concentration (MIC) against representative lab strains (*S. aureus*, *A. baumannii*, and *P. aeruginosa*) - WHO priority pathogens that cause AMR infections in chronic wounds and cystic fibrosis (CF) lungs. MIC data for Bald’s eyesalve is reported in % v/v because it is a liquid preparation. Previous attempts to dry the eyesalve and resuspend the dry mass so MICs could be measured in mg/L reduced its antibacterial activity. We used both antibiotic-susceptible and multidrug-resistant strains in this study. As shown in Table 1, and consistent with previous results (18), Bald’s eyesalve showed good antibacterial efficacy against the majority of the pathogens tested: the MICs for *S. aureus and A. baumannii* were <10% v/v of the reconstructed eyesalve, and are comparable to those of eyesalve batches that were shown to cause a ≥4-log_10_ kill of mature biofilms of *S. aureus* Newman and clinical *A. baumannii* isolates when grown in a synthetic wound model (18). Also consistent with previous results, the MIC for *S. aureus* USA300, an MRSA strain, was higher than its MIC against *S. aureus* Newman, a methicillin-susceptible strain. However, this difference was only a two-fold change in MIC. The highest MIC was seen against *P. aeruginosa* PA14. Interestingly, Bald’s eyesalve showed better inhibitory activity (lower MIC) against a *P. aeruginosa* strain derived from an epidemic cystic fibrosis clone (LESB58) compared to *P. aeruginosa* PA14. Also, Bald’s eyesalve had a slightly better activity (lower MIC) against the multidrug-resistant strain of *A. baumannii* compared to the ATCC 19606 strain.

**Table 1.** Antibacterial activity of Bald’s eyesalve against *S. aureus*, *A. baumannii*, and *P. aeruginosa* in different media. Two strains of *S. aureus* (one MSSA lab strain, Newman, and one MRSA derived from an epidemic pneumonia clone, USA300 LAC) and one lab strain each of *A. baumannii* and *P. aeruginosa* were assayed in the standard antibiotic susceptibility testing medium, caMHB. (Data from three biological replicates (N=3) and three technical replicates (n=3)). MIC is consistent across replicates and in cases of one-fold differences, modal value was taken.

| Strain | Treatment | MIC |
| --- | --- | --- |
| *S. aureus* Newman | Eyesalve (% v/v) | 3.13 |
|  | Vancomycin (µg/ml) | 1 |
| *S. aureus* USA300 | Eyesalve (% v/v) | 6.25 |
|  | Vancomycin (µg/ml) | 1 |
| *A. baumannii* 19606 | Eyesalve (% v/v) | 3.13 |
|  | Meropenem (µg/ml) | 0.5 |
| *A. baumannii* BAA 1605 | Eyesalve (% v/v) | 1.56 |
|  | Meropenem (µg/ml) | 16 |
| *P. aeruginosa* PA14 | Eyesalve (% v/v) | 25 |
|  | Meropenem (µg/ml) | 1 |
| *P. aeruginosa* LESB58 | Eyesalve (% v/v) | 3.13 |
|  | Meropenem (µg/ml) | 16 |

### Bald’s eyesalve causes depolarisation and permeabilisation of the bacterial cytoplasmic membrane, and permeabilisation of the Gram-negative outer membrane

We first assessed whether Bald’s eyesalve had any effect on bacterial membrane integrity or function. The mixture contains bile salts, which cause pore formation in membranes (22), and allicin, which is derived from garlic and onion and is known to interact with membranes and affect their integrity (23). Ethanol and organic acids, typically present in wine, are also known to affect membrane function (24,25). Several assays were carried out on bacteria treated with Bald’s eyesalve for a short period (1-2h). The impact of Bald’s eyesalve on the plasma membrane potential of lab strains of *S. aureus* (MSSA), and *A. baumannii* was assessed with the dye DiSC3(5), which accumulates in the membrane and alters its fluorescence in response to changes in electric potential (26). For both species, we observed an increase in the fluorescence of DiSC3(5) upon the addition of Bald’s eyesalve, which was not seen with the negative control treatment (sterile water) (FIG 1A). This was also similar to the membrane disruption effect seen with 0.1% Triton-X (FIG 1A). This is an indication of membrane disruption. For *A. baumannii*, this membrane disruption occurred both in the presence and absence of polymyxin B nonapeptide (a non-bactericidal membrane permeabiliser) (FIG 1A, Supplementary FIG 1), further confirming the cytoplasmic membrane activity of Bald’s Eyesalve.

Some antibiotics, like polymyxins disrupt bacterial membrane via depolarisation and permeabilisation. To assess if Bald’s eyesalve permeabilises the plasma membrane, we used propidium iodide (PI). PI is a membrane-impermeable nucleic acid dye whose fluorescence increases when bound to the nucleic acid of a membrane-damaged cell. There was increased fluorescence when PI was added to Bald’s eyesalve-treated cells compared to untreated cells, indicating membrane damage in response to Bald’s eyesalve (FIG 1B). An increased number of cells stained with PI (permeabilised cells) was also observed when an established wound biofilm of *S. aureus* was treated with Bald’s eyesalve, further evidencing a bactericidal effect of Bald’s eyesalve via membrane permeabilisation (Supplementary FIG 2).

To further confirm plasma membrane perturbation in response to Bald’s eyesalve, we stained Bald’s eyesalve-treated (at 2×MIC) *E. coli* MG1655 with SYTOX green (nucleic acid stain but only stains permeabilised cells), DAPI (nucleic acid stain), and FM4-64 (membrane stain). This further confirms membrane perturbation, as observed in the SYTOX green and FM4-64 channels (FIG 1C). The membrane perturbation morphology was similar to what was observed with polymyxin B. To further show this membrane perturbation in *S. aureus* and *A. baumannii*, we stained both strains with the plasma membrane stain FM4-64. For *S. aureus*, we observed an increased FM4-64 fluorescence (indicating membrane perturbation) in Bald’s eyesalve- treated cells, which was not seen in the control cells (FIG 1D). With *A. baumannii*, we also observed the presence of dense spots of fluorescence indicative of membrane perturbation (blue arrows) (FIG 1D).

Finally, we assessed if Bald’s eyesalve caused any significant damage to the Gram-negative outer membrane or Gram-positive cell wall by imaging *A. baumannii* and *S. aureus* treated with Bald’s eyesalve with a scanning electron microscope (SEM) (FIG 2). This revealed the presence of morphological changes indicative of cell membrane disruption in Bald’s eyesalve- treated *A. baumannii* cells, which were absent in untreated cells. We did not observe any obvious change in the surface morphology of *S. aureus* under similar treatment, suggesting the cell wall remained largely intact (FIG 2). Negative stain transmission electron microscopy (TEM) was therefore used for *S. aureus* only, and this confirmed that while the cell wall of *S. aureus* was not affected by Bald’s eyesalve treatment, there was observable wrinkling and deformation of the plasma membrane, consistent with the results of the various assays described above (Supplementary FIG 3). For comparison, *S. aureus* treated with antibiotics that either induce membrane pore formation (nisin) or inhibit cell wall synthesis (vancomycin) were also imaged under negative stain TEM; the effect of eyesalve was not visually comparable to that of either antibiotic.

**FIG 1.**
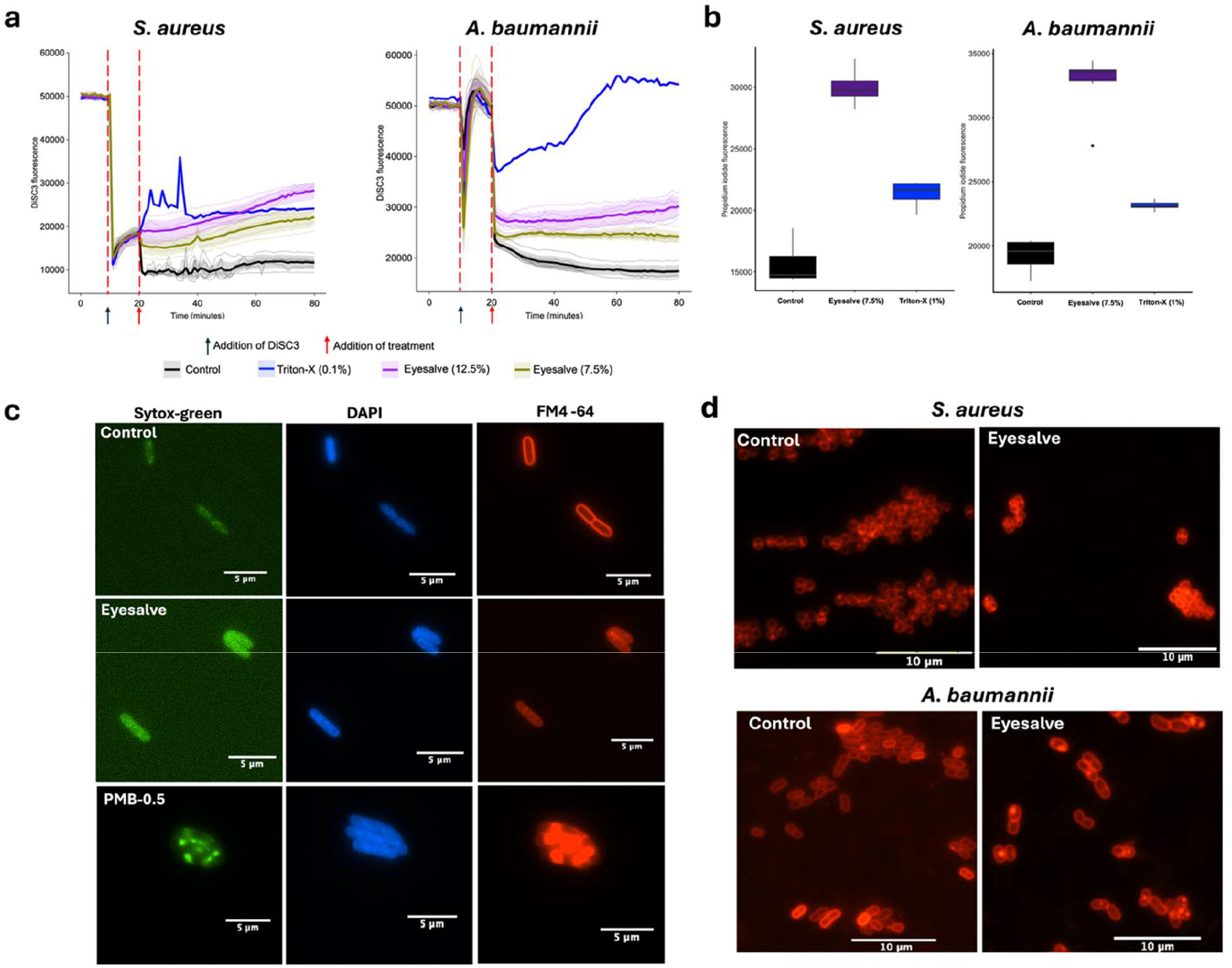
Bald’s eyesalve cause significant disruption of bacterial cytoplasmic membrane. **a)** *S. aureus* Newman and *A. baumannii* ATCC 19606 were stained with DiSC3(5), treated with 12.5% Bald’s eyesalve (purple), 7.5% Bald’s Eyesalve (yellow), 0.1% Triton-X or water (black), and their fluorescence monitored for 1 h. There was increased DiSC3(5) fluorescence, indicating membrane disruption in samples treated with Bald’s eyesalve. The red arrow indicates point of treatment. Results shown are from two biological replicates and three technical replicates; each of the six replicates is shown as an individual line, with the mean and standard deviation plotted as a heavier line and shaded interval. **b)** *S. aureus* Newman and *A. baumannii* ATCC 19606 were treated with 7.5% Bald’s eyesalve (purple), 1% Triton-X or water (grey), stained with propidium iodide, and end fluorescence was measured after 2 h. Box plots show the fluorescence intensity (in RFU) of propidium iodide. Result shown is from two biological replicates and three technical replicates each. Boxes show the interquartile range with median value and whiskers show both minimum and maximum values of the 6 replicates. **c)** *E. coli* MG1655 was treated with 2xMIC eyesalve, 0.5 µg/ml polymyxin B (PMB) or sterile water (control) and stained with SYTOX green, DAPI, and FM4-64. The cells were then imaged with 100x 1.45 NA oil immersion objective on a Nikon TiE widefield epifluorescence microscope. Both eyesalve and PMB showed morphological changes indicative of membrane disruption. Representative images from two replica samples are shown. **d)** *S. aureus* Newman and *A. baumannii* ATCC 19606 were treated with 2×MIC of Bald’s eyesalve or water for 2 h, stained with FM4-64, and then imaged with 100x 1.45 NA oil immersion objective on a Nikon TiE widefield epifluorescence microscope. Increased FM4-64 fluorescence in *S. aureus* and the presence of dense spots of fluorescence in *A. baumannii* suggest membrane perturbation. Representative images from x replica samples are shown.

**FIG 2.**
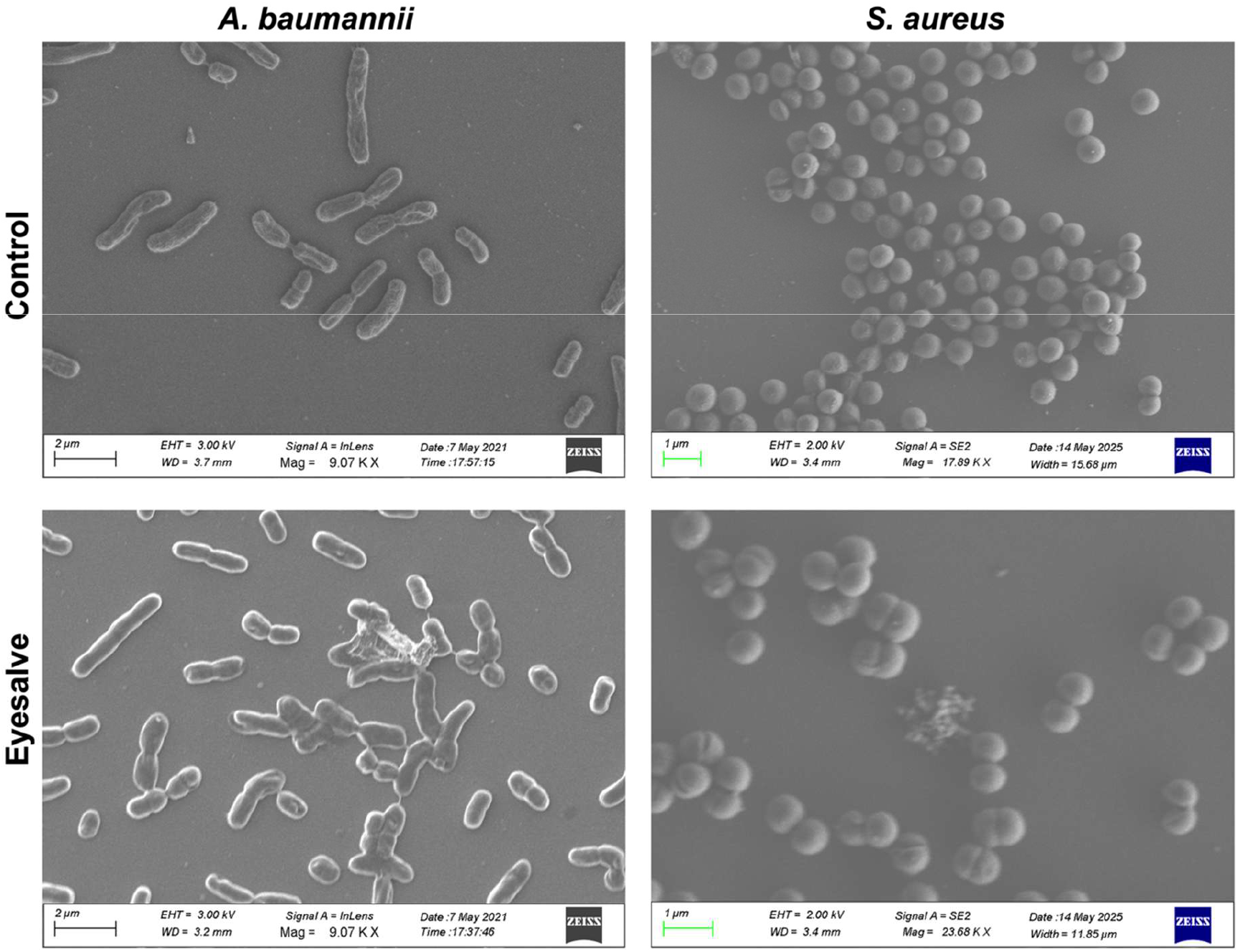
Bald’s eyesalve caused observable deformation of the outer membrane of *A. baumannii*. Both *A. baumannii* ATCC 19606 and *S. aureus* Newman were treated with 2×MIC of Bald’s eyesalve or water for 2 h and imaged with a scanning electron microscope. *A. baumannii* showed signs of disruption in its cell wall/membrane integrity (wrinkling/blebbing) while there were no significant cell wall changes observed in *S. aureus*. Representative images from two replica samples are shown.

### Transcriptomic analysis reveals multiple intracellular responses of *S. aureus* and *A. baumannii* to Bald’s eyesalve, indicating that oxidative stress and DNA damage are part of its mechanism of action

Gene expression analysis using RNA sequencing provides insights into both the primary potential targets of, and downstream responses to, a drug (27,28). To understand gene expression changes in response to Bald’s eyesalve, we carried out RNA sequencing of *S. aureus* and *A. baumannii* treated with 2×MIC of Bald’s eyesalve for 2 h. The cost of RNA sequencing prevented this analysis from being carried out on *P. aeruginosa*. The dose and time point were chosen based on the changes observed with the FM4-64 membrane staining of *S. aureus* and *A. baumannii* treated with 2×MIC of Bald’s eyesalve for 2 h (FIG 1C). Significantly differentially expressed genes were defined as those with p-values < 0.05 and |log2foldchange| ≥1.5 in treated vs. untreated samples., and a full list of these is available in File S1. For *S. aureus*, 346 and 369 genes were significantly up- and downregulated, respectively. For *A. baumannii*, 96 and 68 genes were significantly up- and downregulated, respectively. The functions of these genes were investigated using Gene Ontology (GO) grouping (29). For *S. aureus*, we focussed on the top 100 genes each in the up- and downregulated gene groups in FIG 3 because there were less than 100 genes in each category for *A. baumannii* and GO analysis of all significantly up- and downregulated genes in *S. aureus* yielded too many GO groups. However, we have included all enriched GO groups for *S. aureus* in File S1.

**FIG 3.**
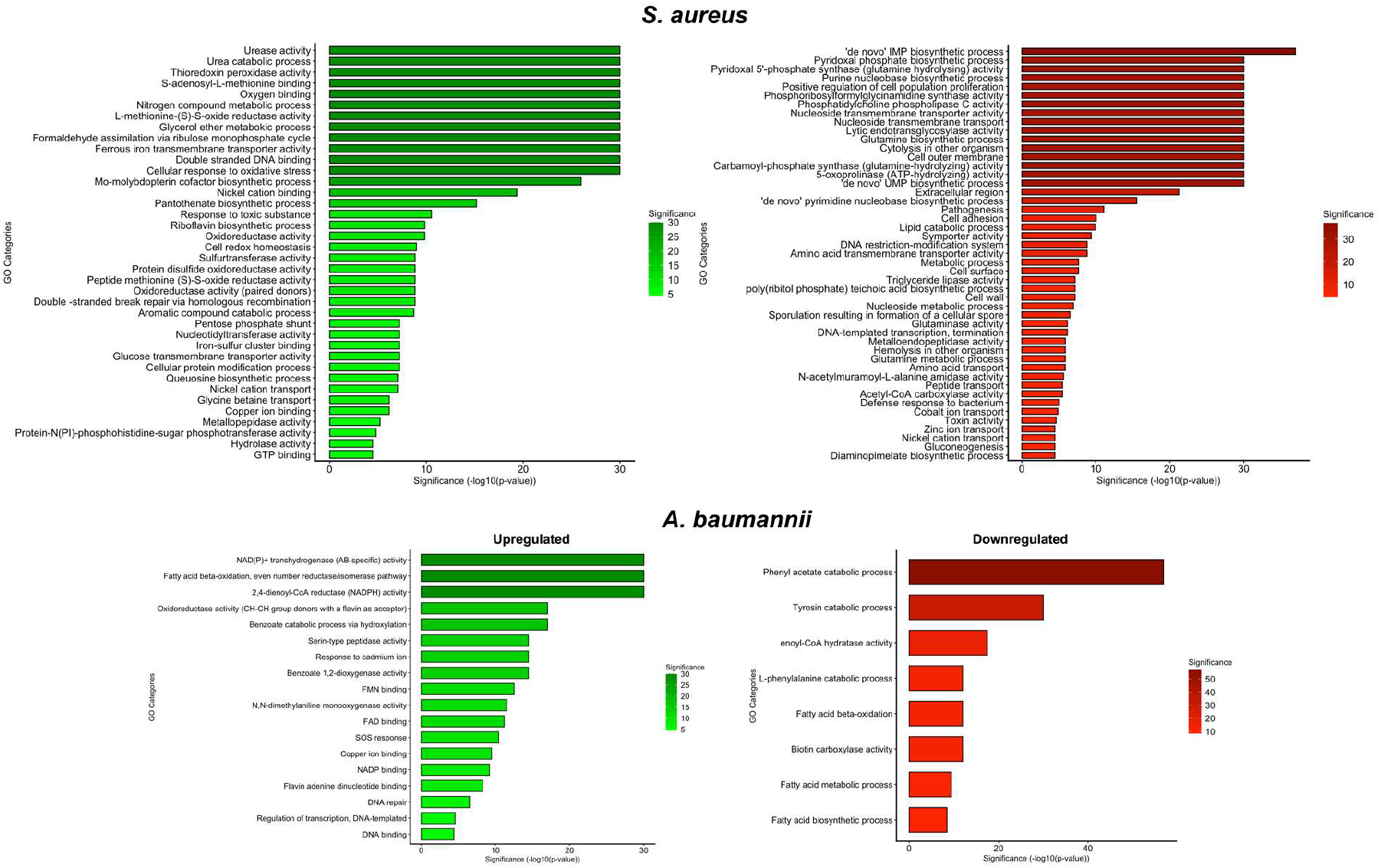
Significantly upregulated and downregulated gene ontology groups in *S. aureus* and *A. baumannii* in response to 2h treatment of 2×MIC of Bald’s eyesalve. Significantly differentially expressed genes were defined as those with p-values ≤ 0.05 and |log2foldchange| ≥1.5 in treated vs. untreated samples; n = 3 and 4 cultures per treatment for *A. baumannii* ATCC 19606 and *S. aureus* Newman, respectively. GO terms are ordered by significance (-log(p-value)).

FIG 3 shows the GO groups of the top 100 significantly differentially upregulated and downregulated genes in *S. aureus*, and all 164 significantly differentially regulated genes in *A. baumannii*. Overall, more GO groups were significantly enriched in *S. aureus* than in *A. baumannii*.

In *S. aureus*, GO groups with significant changes in expression included “cell outer membrane”, “cell surface” and “cell wall”, consistent with the membrane effects shown in phenotypic assays. Membrane-related GO terms were not significantly represented in the response of *A. baumannii,* but upregulation of GO terms associated with cation transport in both species, including “copper ion binding”, “ferrous iron transported activity” and “nickel cation transport/binding”, might be associated with downstream effects of membrane depolarisation.

We also observed significant downregulation of the GO term “pathogenesis”. Notable among these genes are the alpha hemolysin (*hla*), bicomponent leukocidins (*lukGH*), and cysteine histidine amidohydrolase/peptidase (CHAP) domain-containing protein of staphylococcal secretory antigen (*ssaA1*) (File S1). These are important virulence factors produced by *S. aureus* during infections (30,31).

Changes in *S. aureus* expression of “cell redox homeostasis”, “cellular response to oxidative stress”, “oxygen binding”, “oxidoreductase activity” and similar terms appear consistent with the known oxidising ability of allicin (32), which we have shown to be present in the remedy in micromolar amounts (18). This property allows allicin to damage proteins and DNA; DNA damage is also indicated by the upregulation of GOs associated with DNA repair in both *S. aureus* (“double-stranded DNA binding” and “double-stranded DNA repair”) and *A. baumannii* (“DNA binding”, “DNA repair”), and by the upregulation of “SOS response” in *A. baumannii* (File S1).

Gene enrichment analysis also revealed the significant downregulation of genes involved in *de novo* purine and pyrimidine biosynthesis in *S. aureus* (“*de novo* UMP biosynthetic process”, “*de novo* IMP biosynthetic process”, “*de novo* pyrimidine nucleobase biosynthetic process”, and “purine nucleobase biosynthetic process”) (FIG 4). These pathways are essential in the biosynthesis of DNA, RNA, and second messengers, including c-di-GMP (33). The downregulation of genes involved in purine and pyrimidine biosynthesis has previously been linked to increased antibiotic susceptibility and reduced virulence in *S. aureus* (34,35). We also observed the significant downregulation of *pyrP*, which is involved in the pyrimidine salvage pathway, under the “cell outer membrane” GO group.

**FIG 4.**
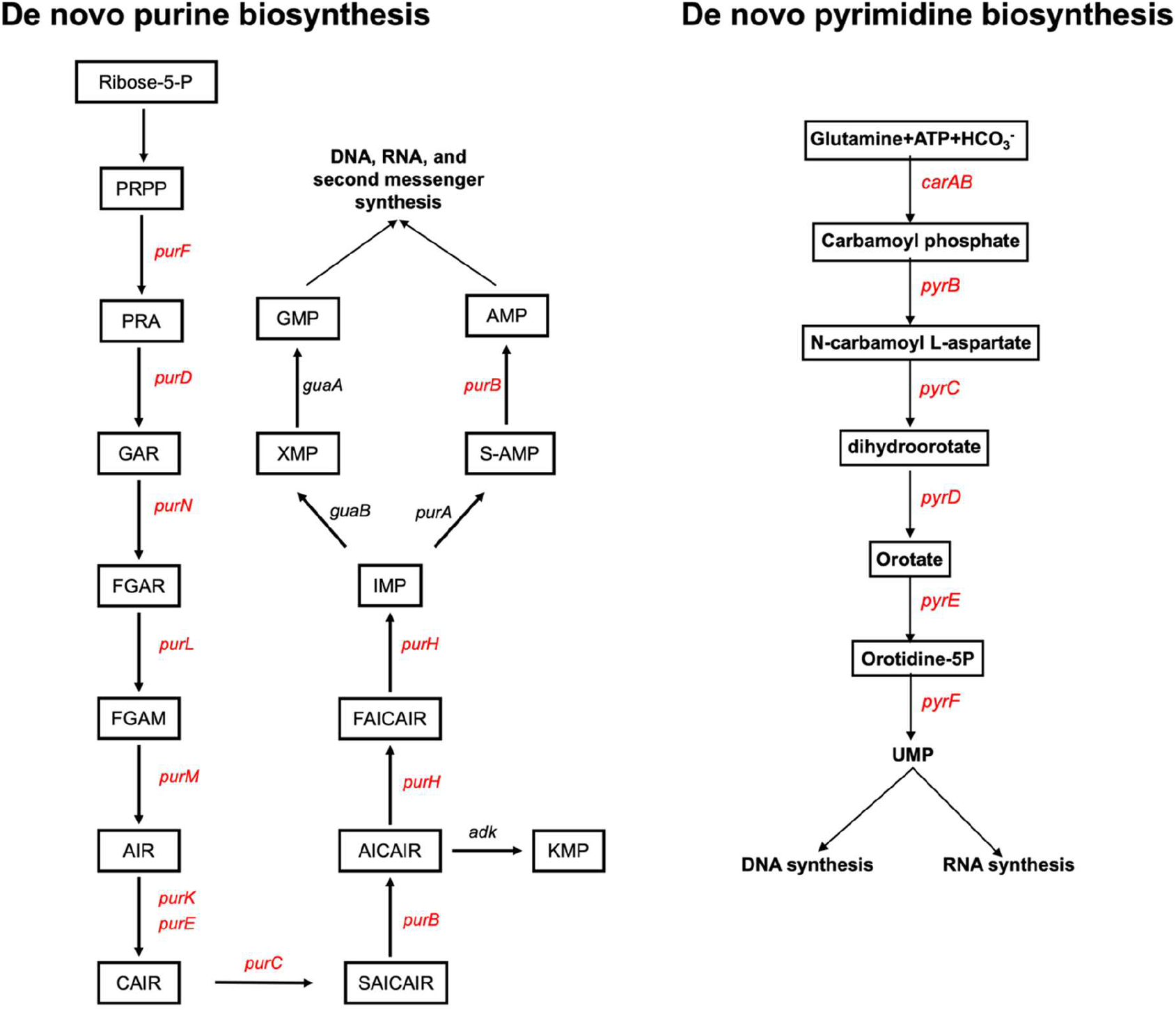
Bald’s eyesalve downregulates the expression of multiple genes involved in *de novo* purine and pyrimidine biosynthesis in *S. aureus* Newman. Representation of the purine and pyrimidine *de novo* biosynthesis pathways in *S. aureus* Newman. Genes in red are significantly downregulated in *S. aureus* Newman in response to Bald’s eyesalve. Significantly downregulated genes were defined as those with p-values ≤ 0.05 and log2foldchange ≤ -1.5 in treated vs. untreated samples; n = 4 cultures per treatment (PRPP = Phosphoribosyl pyrophosphate, PRA = phosphoribosylamine, GAR = glycinamide ribonucleotide, FGAR = phosphoribosylformylglycinamide, FGAM = formylglycinamidine ribonucleotide, AIR = aminoimidazole ribotide, CAIR = carboxyaminoimidazole ribonucleotide, SAICAIR = succinylaminoimidazole carboxamide ribonucleotide, AICAIR = aminoimidazole carboxamide ribonucleotide, FAICAIR = formaminoimidazole carboxamide ribonucleotide, IMP = inosine monophosphate, XMP = xanthosine monophosphate, GMP = guanosine monophosphate, AMP = adenosine monosphosphate, UMP = uridine monophosphate).

In *A. baumannii*, gene enrichment revealed a significant downregulation of GO terms associated with fatty acid metabolism (“fatty acid metabolic process”, “fatty acid biosynthetic process”, “fatty acid beta oxidation”, and “enoyl-CoA hydratase activity”). Fatty acid biosynthesis is important in plasma membrane and lipopolysaccharide biosynthesis in Gram-negative pathogens, and a decrease in its production has been associated with reduced risk of persistence in *A. baumannii* (36). This also aligns with the membrane damage previously reported in FIG 1.

### Bald’s eyesalve inhibits biofilm formation in *S. aureus*

We have previously shown that Bald’s eyesalve can eradicate preformed biofilms of *S. aureus* Newman and *A. baumannii* ATCC 19606 in a soft tissue wound model (18). However, it is not clear if it can also inhibit/prevent biofilm formation at sub-lethal doses in this model. The ability to prevent biofilm formation at low doses would be a very useful property of an antimicrobial. This could help maintain pathogen populations in a more antibiotic-sensitive state in regions of an infection where dosing is limited by poor penetration, or allow administration of low, prophylactic doses to sites vulnerable to infection, for example in applications such as functionalised wound dressings. Genes involved in biofilm formation are divided among several GO terms, including “cell adhesion”, which was significantly downregulated in *S. aureus* (Figure 3). To explore the potential of Bald’s eyesalve to prevent biofilm formation, we therefore manually searched our dataset of DEGs to identify genes involved in biofilm formation.

In *S. aureus*, some genes involved in biofilm formation were significantly differentially expressed and grouped under the GO term for “cell adhesion” including *sdrC, sdrD, sdrE,* and *cflB* (Supplementary FIG 4). Overall, only genes involved in the attachment stage of biofilm formation in *S. aureus* were differentially downregulated (Supplementary FIG 5). The majority of the biofilm-associated genes in *A. baumannii,* including genes responsible for the production of the biofilm exopolysaccharide, poly-N-acetyl glucosamine (PNAG) (*pgaABCD*), pili - for attachment (*csu* genes), and biofilm-associated proteins in *A. baumannii* (*bap*), were significantly downregulated (Supplementary FIG 4). Because Bald’s eyesalve caused downregulation of loci involved in biofilm formation, we used the optotracer EbbaBiolight 680 to assess the phenotypic effect of sub-bactericidal eyesalve treatment on production of biofilm extracellular polymeric substances. EbbaBiolight 680 binds bacterial amyloids and glucans, and fluoresces when bound to these molecules; it has successfully been used to assess production of biofilm matrix in Gram-positive and Gram-negative bacteria, including *S. aureus* (37,38). We assessed the effect of treatment with 1× MIC eyesalve (3.125% v/v) on EbbaBiolight 680 fluorescence over 24h when it was spiked into planktonic cultures of *S. aureus* Newman and *A. baumannii* initiated by diluting stationary-phase bacteria into fresh caMHB. The optotracer did not produce any measurable fluorescence in the absence of bacteria, regardless of whether eyesalve was added to the media (Supplementary FIG 5).

As shown in FIG 5A, eyesalve treatment caused a slight decrease in the log-phase growth rate and stationary phase density of *S. aureus*, but ablated the steady increase in optotracer fluorescence observed during stationary phase of untreated cultures. This is consistent with the eyesalve specifically reducing production of biofilm exopolymers at this dose. However, as shown in FIG 5B, despite the slight decrease in log-phase growth rate and delayed entry into stationary phase caused by eyesalve treatment of *A. baumannii*, eyesalve treatment expedited and increased optotracer fluorescence, consistent with earlier and greater production of biofilm exopolymers at this dose. It is well known that bacteria sometimes upregulate biofilm production at certain sub-inhibitory doses of antibiotics and antimicrobials (39,40), and a finer-grained exploration of dose dependence of bacterial responses would be necessary to determine the relationship between positive and negative effects on biofilm formation, and eyesalve dose. With the available data, we have good phenotypic confirmation that sub-bactericidal doses of Bald’s eyesalve can inhibit biofilm formation in *S. aureus*, but not in *A. baumannii*.

### Bald’s eyesalve represses quorum sensing in *S. aureus*

Due to the inhibitory effect of Bald’s eyesalve on the expression of *S. aureus* genes encoding acute virulence factors in the GO group “pathogenesis”, we decided to specifically explore the expression profile of *agr* genes in *S. aureus*. The *S. aureus* Agr quorum-sensing system plays a significant role in controlling biofilm formation and virulence factor expression (FIG 6A). Transcriptomic analysis revealed the downregulation of *agrA*, *agrC*, and *agrB* in response to Bald’s eyesalve (FIG 6A). AgrB processes a precursor of the auto-inducing peptide (AIP) signal, encoded by *agrD*, and exports the mature signal from the cell. AgrC and AgrA form a two-component system that senses extracellular AIP and, once the quorum is reached, activates transcription of both the *agrACDB* operon and of RNAIII, which in turn regulates loci controlling biofilm formation and virulence factor expression (FIG 6A).

To phenotypically confirm the repression of Agr-mediated quorum sensing by Bald’s eyesalve, we assessed the effect of sub-inhibitory concentrations of Bald’s eyesalve on two bioreporter strains: a constitutive reporter encoding GFP and a lux cassette under the control of a growth- dependent promoter (P*xylA*:gfp:lux), and an AIP-dependent reporter containing the same construct under the control of the RNAIII promoter (P3:*gfp*:l*uxCDABE*) (41). We observed concentration-dependent quorum-sensing inhibition of the AIP-dependent reporter at low, sub- inhibitory concentrations of Bald’s eyesalve (0.33% and 0.033%), which was not observed in the growth-dependent reporter (FIG 6B, Supplementary FIG 6). A slightly higher sub- inhibitory concentration (3.3%) caused a reduction in reporter expression from both promoters, consistent with this concentration causing general cellular stress as outlined above. Quorum- sensing reporters were not available for *A. baumannii*, but they are available for the other Gram-negative species explored phenotypically in this study, *P. aeruginosa*. In an experiment analogous to that described for *S. aureus*, we found that Bald’s eyesalve did not affect the *las* (3-oxo-C12 homoserine lactone-mediated) quorum-sensing system, which positively regulates acute virulence in this species (Supplementary FIG 7).

### Laboratory selection for resistance to Bald’s eyesalve led to slower resistance evolution, compared with mainline antibiotics

Resistance evolution is a major challenge in clinical settings, and because various models have predicted that resistance evolution may be harder to achieve for antimicrobials with multiple targets (42,43), we wanted to check how rapidly or slowly pathogens would evolve resistance to Bald’s eyesalve under strong selection for resistance. To achieve this, we carried out a 14- day selection experiment using an evolutionary ramp approach. We initially grew planktonic populations (three populations per bacterial species) of *S. aureus* Newman, *A. baumannii* ATCC 19606, and *P. aeruginosa* PA14 for 24 h in 0.5×MIC of Bald’s eyesalve or an example clinically-relevant antibiotic, and then passaged replica aliquots of each population to fresh medium containing either 0.5× or 1×MIC. If a population grew at the higher of the two concentrations, it was passaged to fresh media with 1×MIC and 2×MIC of the treatment; if it failed to grow in the higher concentration, it was passaged to 0X or 0.5×MIC. In this way, populations that continued to grow were challenged with higher concentrations of treatment, increasing by two-fold each passage after successful growth. The results are shown in (FIG 7) and (Table 2). At the end of day 14, the bacteria were growing in 2-256×MIC of antibiotic, but only 0.5-2×MIC of Bald’s eyesalve. Thus, we observed much slower resistance evolution to Bald’s eyesalve, suggesting that it is more difficult for bacteria to evolve resistance to this complex cocktail than to single-molecule antibiotics with a narrower range of cellular targets. We also attempted to select for resistant *S. aureus* by serially passaging in a constant sub- inhibitory concentration of ESO for 28 daily transfers. This also failed to generate any resistant mutants (data not shown).

**FIG 5.**
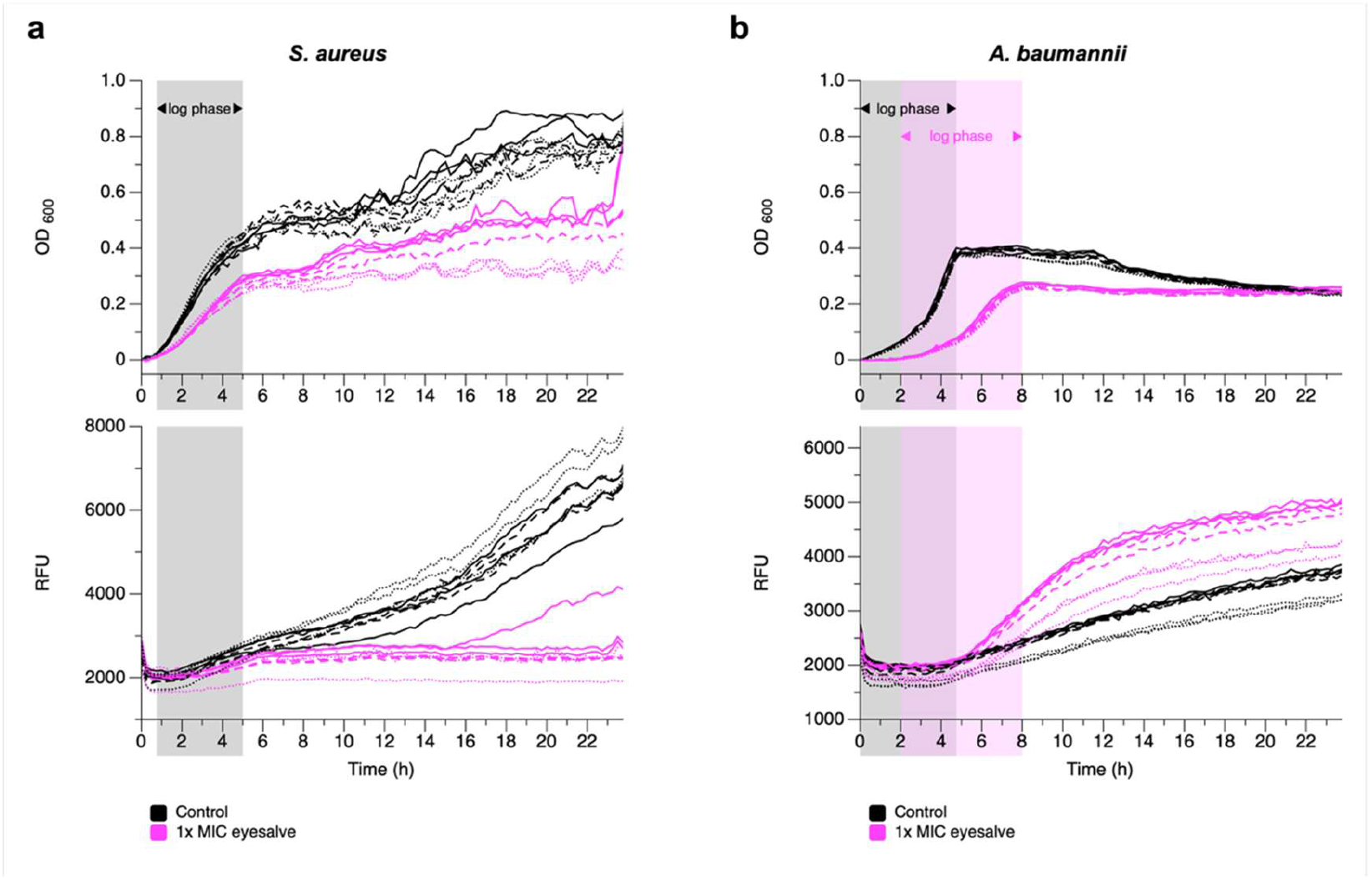
Bald’s eyesalve inhibits production of biofilm matrix exopolymers by *S. aureus*, but not by *A. baumannii*. Stationary-phase cultures of *S. aureus* Newman (A) and *A. baumannii* ATCC 19606 (B) at 0.08-0.1 OD600 were diluted into fresh caMHB and exposed to Bald’s eyesalve at 1× MIC eyesalve (3.125% v/v, pink lines), or left untreated (black lines), in the presence of 1 µg/ml of EbbaBiolight 680 for 24 h. EbbaBiolight 680 is an optotracer which fluoresces when bound to biofilm exopolymers. Three aliquots each of three independent overnight starter cultures were used in this experiment: each line represents one experimental culture, with the three starter cultures denotes by solid, dashed and dotted lines. Top panels show OD600, standardised to show the initial OD as zero. Both untreated and treated *S. aureus* had similar onset and end times for log phase, which is shown by the grey shading. Treated and untreated *A. baumannii* had noticeably different onset and end times for log phase, shown by the grey and pink shading, respectively. Bottom panels show relative fluorescence units (RFU) at ex/em 550/670 nm. Log phase shading has been added to these panels for each of comparison with the OD data above, and to show that RFU dynamics over time do not simply reflect cell density.

**FIG 6.**
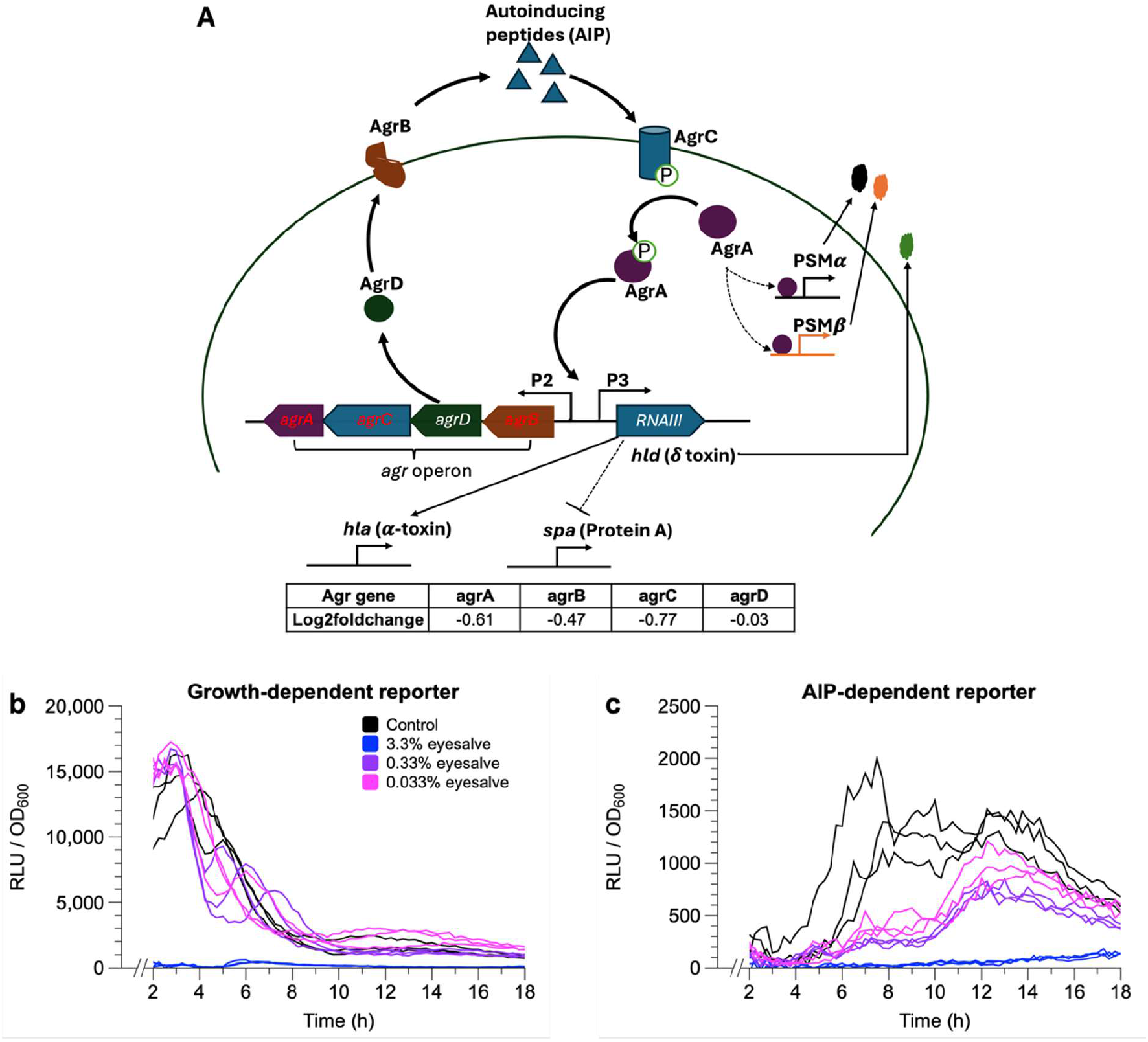
Bald’s eyesalve inhibits quorum-sensing signalling in *S. aureus*. A. Downregulation of the expression of *agrA*, *agrB*, and *agrC,* which are involved in both quorum-sensing and virulence factor production in *S. aureus*. B and C. Mid-log phase cultures at 0.08-0.1 OD600 of *S. aureus* RN6390 P*xylA*:*gfp*:*luxCDABE* (a growth-dependent reporter/constitutive promoter), *S. aureus* RN6390 P3:*gfp*:l*uxCDABE* (autoinducing peptide (AIP)-dependent reporter/QS-dependent promoter) were exposed to Bald’s eyesalve concentrations of 3.3%, 0.33% or 0.033% v/v concentrations of Bald’s eyesalve, or to sterile water, for 18 h and OD600 and luminescence (relative light units, RLU) measured at intervals to assess the effect on quorum sensing. Each line represents an individual bacterial culture. The highest concentration of Bald’s eyesalve used approximates the MIC against *S. aureus* Newman in caMHB (Table 1), but because this assay uses a starting inoculum of bacteria that is more dense than that used in MIC assays, and also at mid-log phase, this concentration does not affect growth (Supplementary FIG 7). Plotting RLU/OD from the start of log growth phase shows that for both reporters, 3.3% eyesalve completely prevented luminescence. For the growth-dependent reporter, lower concentrations had no discernible effect on expression. However, there was concentration-dependent repression of the reporter gene in the AIP-dependent reporter, shown by the peak of luminescence becoming progressively lower and later with addition of 0.033% and 0.33% eyesalve. This is consistent with 3.3% eyesalve causing generalised cellular stress, but lower concentrations specifically affecting AIP-mediated quorum sensing

**FIG 7.**
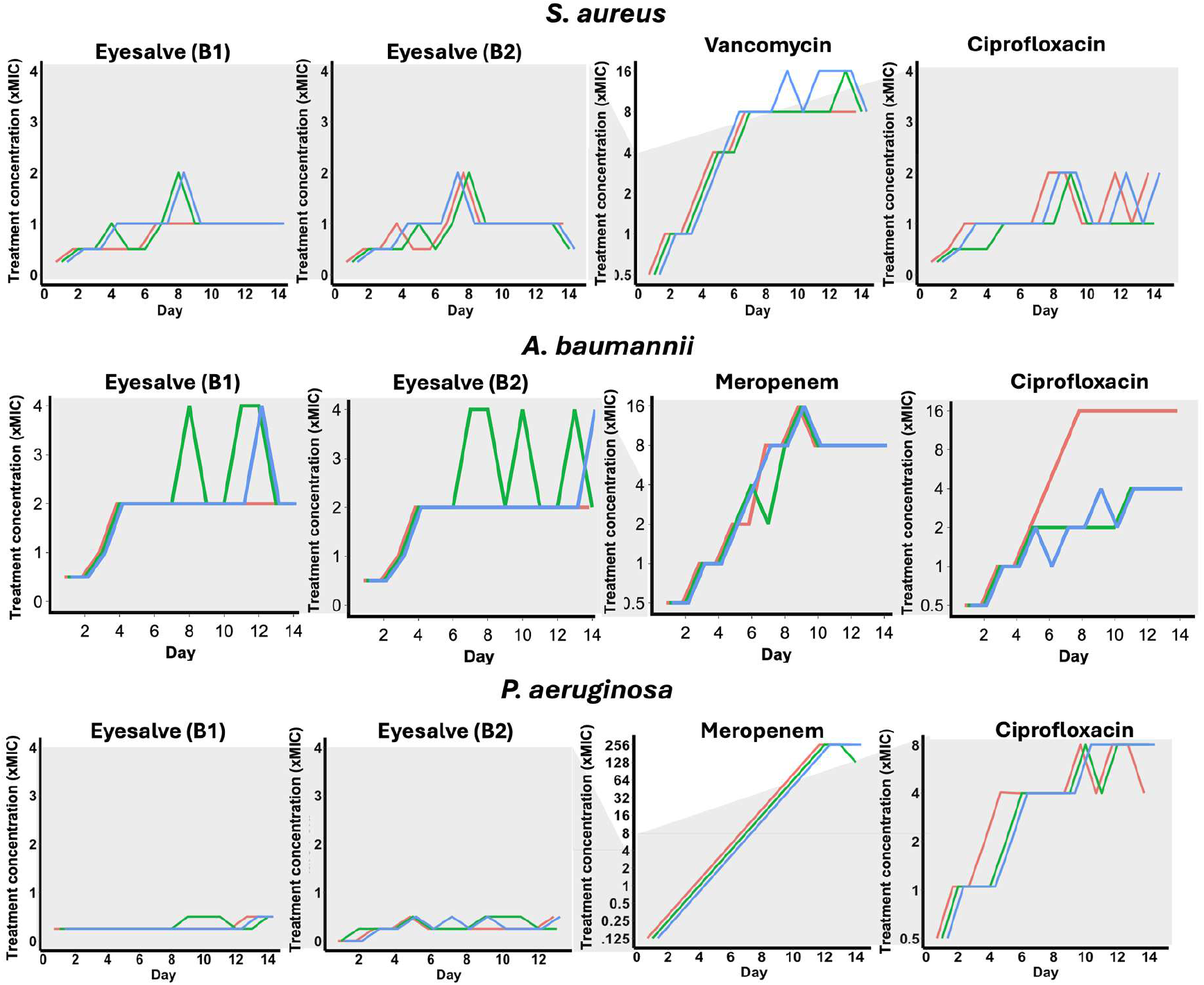
Slower resistance evolution to Bald’s eyesalve compared to mainline antibiotics. *S. aureus* Newman, *A. baumannii* ATCC 19606, and *P. aeruginosa* PA14 were subjected to subinhibitory concentrations of Bald’s eyesalve or antibiotic and subsequently passaged in different concentrations based on the previous day’s highest concentration with growth. This was done for 14 days. The graph’s axes show the highest of two tested concentrations in which cultures grew at each time point, expressed as a multiple of the ancestral MIC. Each line shows one of three independent populations.

**Table 2.** Change in the concentration of Bald’s eyesalve in which *S. aureus* Newman, *A. baumannii* ATCC19606, and *P. aeruginosa* PA14 could grow after 14 days of selection in an evolutionary ramp experiment.

| Bacteria | Treatment | Fold change in growth concentration at day 14 |
| --- | --- | --- |
| *S. aureus* Newman | Vancomycin | 8 x MIC |
|  | Ciprofloxacin | 2 x MIC |
|  | Eyesalve (batch A) | 1 x MIC |
|  | Eyesalve (batch B) | 0.5 x MIC |
| *A. baumannii* ATCC19606 | Meropenem | 8 x MIC |
|  | Ciprofloxacin | 16 x MIC |
|  | Eyesalve (batch A) | 2 x MIC |
|  | Eyesalve (batch B) | 4 x MIC |
| *P. aeruginosa* PA14 | Meropenem | 256 x MIC |
|  | Ciprofloxacin | 8 x MIC |
|  | Eyesalve (batch A) | 0.5 x MIC |
|  | Eyesalve (batch B) | 0.5 x MIC |

## DISCUSSION

The increasing mortality and economic cost associated with AMR have necessitated the need for new antimicrobials (1,2). It is even more important to have new antimicrobials with multiple modes of action to reduce the risk of resistance. We previously identified a historical remedy called Bald’s eyesalve and made a reconstruction of this remedy that had broad-spectrum antibacterial and antibiofilm efficacy against both Gram-positive and Gram-negative pathogens (17–19). We also showed that this reconstructed historical remedy has great antibacterial activity against *S. aureus*, *S. epidermidis*, *Streptococcus pyogenes*, *Enterococcus cloacae*, *A. baumannii*, and *P. aeruginosa* (18). Safety analysis of Bald’s eyesalve has so far shown that it has little or no adverse effect in *in vitro* and *in vivo* models (19,20).

In this study, we have shown that Bald’s Eyesalve has good (<10% v/v of the reconstructed Eyesalve inhibited bacterial growth) antibacterial effect on *S. aureus*, *A. baumannii*, and *P. aeruginosa,* including multidrug resistant strains (MRSA and carbapenem-resistant *A. baumannii* and *P. aeruginosa*). The antibacterial effect of this historical remedy is multifaceted, targeting both the cell membrane(s) and intracellular activities. Bald’s eyesalve perturbed the membrane potential of both Gram-positive (*S. aureus*) and Gram-negative (*A. baumannii* and *P. aeruginosa*) pathogens. Bacterial membrane potential controls important cellular physiological processes in cells, including membrane transport, motility, ATP synthesis, and cell division (44). This also correlated with the significant downregulation of genes encoding membrane-associated transport proteins, including some efflux pump proteins like NorB (*S. aureus*) and AdeT1 (*A. baumannii*). NorB and AdeT1 belong to the major facilitator superfamily (MFS) and resistance nodulation division (RND) classes of efflux pumps, respectively, and confer multidrug resistance on these pathogens (45). The downregulation of these genes could signify the inhibition of the antimicrobial resistance machineries of these pathogens, and future work could usefully seek to phenotypically validate this prediction.

Furthermore, there was downregulation of biofilm- and virulence-associated genes in both *S. aureus* and *A. baumannii* - most especially, cell surface attachment proteins. The first and essential stage of biofilm formation involves surface or non-surface attachments (46). This attachment stage allows bacterial pathogens to form microcolonies that develop into mature biofilms; hence, an inhibition of this process prevents biofilm formation. Cell surface attachment proteins are also essential for host cell attachment during infections (47). In addition to inhibiting attachment, Bald’s eyesalve also inhibited expression of biofilm exopolysaccharide and protein-associated genes in *A. baumannii*. Consistent with downregulation of biofilm- and virulence-associated genes, Bald’s eyesalve caused downregulation of *S. aureus* genes involved in quorum sensing, and a corresponding reduction in AIP production was phenotypically validated using fluorescent bioreporters. It is possible that some of the other gene expression patterns in *S. aureus* might not be a direct mechanism of Bald’s eyesalve but a downstream effect of its effect on quorum sensing.

Furthermore, Bald’s eyesalve significantly downregulated fatty acid metabolism in *A. baumannii*. Increased fatty acid production has been associated with antibiotic persistence in *A. baumannii* (36). Hence, the significant downregulation of this process might correlate with increased antibiotic susceptibility in *A. baumannii* to Bald’s eyesalve. Also, due to the role of fatty acids in the biosynthesis of phospholipids and lipopolysaccharides in Gram-negative pathogens, inhibition of fatty acid biosynthesis has been suggested as a good target mechanism for new antimicrobials (48). This is another of the transcriptomic responses which we were unable to phenotypically validate due to time and cost, and which could be explored in future.

This multifaceted mechanism of action of Bald’s eyesalve is likely linked to the combinatorial effect of all its component ingredients. Each of the components (onion, garlic, wine, and bile) of this historical remedy has been reported to have some antibacterial effect. Both garlic (*A. sativum*) and onion (*A. cepa*), the two most popular members of the *Allium* genus and some of the world’s oldest crops, are well known for their medicinal benefits (49,50). These aromatic spices, most especially garlic, have been reported for their bioactivities, including anticancer (51), antioxidant (52), antifungal (53), and antibacterial activities (54,55). The antibacterial activity of garlic has been reported against a broad range of both Gram-positive and Gram-negative pathogens *in vitro* (54,56). Some of the common bioactive compounds in garlic and some other *Allium* species include allicin, ajoene, phenols, flavonoids, saponins, triterpenes, steroids, and other organic acids (49,57). The most popular and most studied for its antibacterial efficacy, among them, is allicin (23,32). We have previously shown that the antibacterial efficacy of Bald’s eyesalve is not just due to the presence of allicin (18). Wine, the third component of Bald’s eyesalve, is one of the oldest and most popular alcoholic beverages in the world and is mostly produced from the fermentation of grape must or juice (58). In addition to the popular use of wines for celebrations, they have also been known for centuries for their health benefits due to their variety of bioactive compounds (59,60). Although the optimal antimicrobial activity of wine is majorly due to the combinatorial effect of all its components (pH, alcohol, phenolics, and organic acids) (61), the majority of its antimicrobial effect is associated with the different phenolics and organic acids in it (60,62,63). Bile, the last component of Bald’s eyesalve, have been previously shown to have inhibitory activity against bacterial strains, including *S. aureus* and *P. aeruginosa,* as well as prevent biofilm formation in these strains (64,65). The ability of Bald’s eyesalve to inhibit *S. aureus* biofilm formation was also evidenced in this study. The antibacterial activity of bile salts has been associated with the disruption of bacterial membranes, leading to the release of intracellular content (64).

Expectedly, this multifaceted mechanism of action of Bald’s eyesalve reduced the risk of resistance evolution against it compared to clinically relevant antibiotics. This is in line with previous studies that reported slow resistance evolution to antimicrobials with multiple targets (42,43). Additionally, the slower resistance evolution seen with Bald’s eyesalve might be associated with its dual effect (depolarisation and permeabilisation) on bacterial membrane (66). Again, impaired resistance evolution by *S. aureus, A. baumannii*, and *P. aeruginosa* against Bald’s eyesalve can be linked to the combinatorial effect of its multiple components which target different parts of these bacterial pathogens as shown in this study.

Taking together previous results on Bald’s eyesalve’s antibacterial and antibiofilm efficacy and the mechanism of action shown in this study, we conclude that this historical remedy has great potential to be used as a foundation for developing an antimicrobial for bacterial infections, especially those resulting from *S. aureus* and *A. baumannii*. We still face challenges associated with its biofilm eradication activity against *P. aeruginosa* and batch-to-batch variability (18). Future studies on Bald’s eyesalve should focus on (1) identifying the specific compounds present that confer antibacterial, antibiofilm and/or antivirulence activity and creating candidate synthetic molecular cocktails for activity testing; and (2) identifying synergists or potentiators that improve the ability of Bald’s eyesalve to eradicate *P. aeruginosa* chronic wound biofilm and reduce its batch-to-batch variability. Our end goal is not proposing the use of Bald’s eyesalve in the clinic, but to gather sufficient information to develop a more defined and clinically relevant formulation of natural products based on this complex remedy, its effects on bacteria and its interacitons with other antimicrobials. This will be the focus of future studies on this preparation.

## MATERIALS AND METHODS

### Bacterial strains

The bacterial strains used in this study are *S. aureus* Newman, *S. aureus* USA300 LAC, *P. aeruginosa* PA14, *A. baumannii* ATCC 19606, and BAA 1605, *S. aureus* RN6390 P*_xylA_*:*gfp*:*luxCDABE* (a growth-dependent reporter), *S. aureus* RN6390 P3:*gfp*:l*uxCDABE* (autoinducing peptide (AIP)-dependent reporter) (41), *P. aeruginosa* NPAO1 (Nottingham PAO1 lab strain) mini-CTX::*luxCDABE* (growth-dependent reporter), *P. aeruginosa* NPAO1 mini-CTX::*lasB-luxCDABE* (C12-homoserine lactone-dependent reporter), and *Escherichia coli* MG1655.

### Media

The bacteriological media used in this study were phosphate buffered saline (PBS), cation-adjusted Mueller Hinton broth (caMHB) (Sigma-Aldrich, USA), synthetic wound fluid (SWF) (50% v/v foetal bovine serum (Gibco) and 50% v/v peptone water (Sigma-Aldrich, USA) (67) and Luria Broth (LB) agar (Sigma-Aldrich, USA).

### Bald’s eyesalve preparation

Bald’s eyesalve was prepared as previously reported (17,18). Briefly, equal volumes of onion and garlic sourced locally were chopped separately into pieces with a sterile knife and then pounded together with a sterile mortar and pestle for 2 min. Equal amounts of wine (Pennard Organic, Somerset, UK) and oxgall (Sigma Aldrich, USA) were then added, and the mixture was placed in a sterile 250 ml Duran bottle covered with aluminium foil. The bottle was placed in the fridge (at 4°C) for 9 days. After 9 days, the lumps in the bottle were strained and the liquid transferred into sterile 50 ml Falcon tubes. The tubes were centrifuged at 3000 rpm for 5 minutes, after which the liquid was filtered into 2 ml Eppendorf tubes with a sterile syringe and a 5 µm filter. Samples were stored in -20°C for further analysis.

### Minimum inhibitory concentration assay

The broth microdilution method was used to determine the minimum inhibitory concentration of the antimicrobials as recommended by the European Committee on Antimicrobial Susceptibility Testing (EUCAST). Briefly, the bacterial strain of interest was streaked on LB agar and incubated at 37°C for 18-24 h to produce distinct colonies. Twice the maximum concentration of the antibiotics to be used was prepared in cation-adjusted Müller-Hinton broth (caMHB) and dispensed (100 µl) in the first column of a Corning Costar CLS9018 (Corning Inc., US) 96-well plate. Fifty microlitres of the medium were then dispensed into each of the other wells, after which a two-fold serial dilution was carried out. A bacterial suspension in the medium was prepared by touching 3-4 distinct colonies with a sterile cotton swab and dispersing them in PBS. This was then standardised to a 0.5 MacFarland standard with a spectrophotometer (OD at 600nm = 0.08-0.10) and further diluted to a starting concentration of 1x10^5^ CFU. The standardised bacterial suspension (50 µl) was then inoculated into triplicate wells of each concentration except for the sterile control well (without antimicrobials and bacteria). Growth control wells (with only bacteria) were also set up. The 96-well plates were sealed with Parafilm^TM^ and incubated at 37°C for 18-24 h. The lowest concentration well with no observable growth is taken as the MIC. Uninoculated medium (without treatment) was used as a negative control, while bacterial suspension without treatment was used as a positive control. Meropenem and vancomycin were used as antibiotic controls for Gram-positive and Gram-negative bacteria, respectively.

### DiSC3(5) kinetics assay for cell membrane disruption

A DiSC3(5) (3,3’-Dipropylthiadicarbocyanine iodide) assay was performed as previously described by Buttress et al. (26) with a little modification. Briefly, bacterial culture growing at the log phase in LB was diluted to 0.5 OD_600_ with PBS containing 0.5 mg/ml bovine serum albumin (BSA) (Sigma-Aldrich, UK) and 2 µM glucose. The addition of BSA was to prevent the binding of DiSC3(5) reagent to the microtitre plate. Bacteria cells (*S. aureus* and *A. baumannii*) were washed in PBS containing 0.5 mg/ml bovine serum albumin (BSA) (Sigma-Aldrich, UK) and 2 µM glucose twice and then resuspended. After resuspension, the tube containing washed cells was incubated for 15 min. Bacteria were then transferred into a black polystyrene 96-well plate (Corning Inc., US) and autofluorescence measured (excitation wavelength = 610±10, emission wavelength = 660±10) for 10 min in Tecan SPARK 10 M (Tecan, Switzerland) plate reader, after which DiSC3(5) reagent (Invitrogen) in 1% DMSO was added to give a final DiSC3(5) concentration of 0.5 µM and fluorescence measured for another 10 minutes. Treatments were then added, and the fluorescence was monitored for 1 h. 0.1% Triton-X was used as a positive control.

### Cytological profiling

Overnight cultures of *E. coli* were grown to mid-log phase and diluted to 0.1-0.2 0.1-0.2 OD_600_ in LB. The diluted cultures were then treated with media or 2xMIC Eyesalve for 1 h. Cells were then washed by centrifuging at 10,000 rpm for 5 min and resuspended in PBS. Cells were stained with FM4-64 (red), DAPI (blue) and Sytox Green (green). The stains were then washed, and cells were fixed with formaldehyde. The images were taken with 100x 1.45 NA oil immersion objective on a Nikon TiE widefield epifluorescence microscope. Scale bar equivalent of 10µm. For *S. aureus* and *A. baumannii*, overnight cultures of bacteria were grown to mid-log phase and diluted to 0.1-0.2 OD_600_ in caMHB. The diluted cultures were then treated with media, 2×MIC of Bald’s eyesalve for 2 h. Cells were then washed by centrifuging at 10,000 rpm for 5 min and resuspended in PBS. FM4-64 was then added to a final concentration of 2.5 µg/ml and incubated for 15-20 min. The stain was then washed, and cells were fixed with formaldehyde. The images were taken with a Nikon Eclipse-Ti wide-field fluorescence microscope at oil immersion magnification.

### Live-dead assay with propidium iodide

A membrane permeabilisation assay was carried out as previously described by Boix- Lemonche (2020), with modifications. First, bacteria at mid-log phase of growth in LB were diluted to an 0.1 OD_600_ with PBS containing 2 µM glucose, transferred (50 µl) into a black 96-well flat bottom plate CLS3596 (Corning Inc., US) and autofluorescence (excitation wavelength = 530±20, emission wavelength = 620±10) was measured for 5 min in Tecan SPARK 10 M (Tecan, Switzerland) plate reader. Propidium iodide (50 µl) (Invitrogen) was added to the wells at a final concentration of 50 µg/ml, followed by fluorescence measurement for another 5 min. Treatments (sterile water (negative control), 7.5% Eyesalve, 1% TritonX (positive control)) were then added, after which fluorescence was measured for 1 h. Live-dead staining of *S. aureus* biofilm was done with LIVE/DEAD BacLight staining reagent (Invitrogen) - containing propidium iodide and SYTO 9 - as described by the manufacturer, with modifications. A thin film of collagen wound matrix was made by pipetting 200 µl of wound matrix (prepared as previously described) into a 24-well insert (VWR International, UK) on a sterile glass microscope slide placed in a Petri dish and UV sterilised as previously described. This was infected with bacterial suspension (50 µl) at log phase of growth, diluted to 0.08-0.1 OD_600_ (approximately 10^8^ CFU). The Petri dish was covered and sealed with Parafilm^TM^ to prevent it from drying out. This was then incubated at 37°C for 24 hours to allow biofilm formation. The formed biofilms were then treated with either PBS, Bald’s eyesalve and incubated overnight. The slides were then stained with a mixture of components A and B from the LIVE/DEAD BacLight staining kit and incubated for 15 minutes. The slides were then fixed with 4% formaldehyde, washed three times with PBS, and viewed using a confocal laser scanning microscope (Zeiss LSM 880) with the oil immersion magnification (x100) lens.

### Scanning electron microscopy

Overnight bacterial culture was resuspended in caMHB to 0.1 OD_600_ and incubated on a shaker at 37°C until it reached mid-log phase (10^6^ to 10^7^ CFU/ml). Cultures were then diluted (equal volume) with treatment (Bald’s eyesalve, or caMHB (control)) and incubated for 2 h. Bacterial cells were then centrifuged at 10000 rpm for 1 min, followed by 3 washes of PBS. Twelve (12) millimetre coverslips (VWR, US) were then incubated with 50 µl of poly-lysine (Sigma-Aldrich, US) for 15 min, after which the poly-lysine was removed and the coverslips left to dry. Bacterial cell pellets were resuspended in 400 µl of PBS, after which 50 µl of this was added to each cover slip (in a 24-well plate) and incubated for 30 min at 37°C. Excess (unbound) bacterial suspension was then discarded, and the bound cells were fixed with 2.5% glutaraldehyde solution in PBS overnight. After fixing, excess glyceraldehyde was removed, and the coverslips were washed 3 times with PBS. The coverslips were transferred into clean wells and dehydrated using ethanol gradient (20%, 50%, 70%, 90%, 100%, and 100%) for 10 min in each ethanol concentration. The coverslips were then transferred into new wells and incubated with 0.5 ml of hexamethyldisilazane (HDMS) for 30 min. The coverslips were then moved again into new wells, the HDMS discarded, and the slips were left to dry in a laminar flow cabinet for 30 min. Coverslips were then added to copper tapes placed on SEM sample holders, and the samples were sputtered with carbon. The images were taken with the Zeiss Gemini Scanning Electron Microscope at the Warwick Electron Microscope Research Technology Platform.

### Negative staining and transmission electron microscopy

*S. aureus* Newman was grown in SWF for 6h to allow cells to reach the log phase in a 96- well plate. In the wells, 50 µl of treatment (water, nisin – 150 µg/ml, vancomycin – 100 µg/ml, and 100% Bald’s eyesalve) was added to 100 µl of bacterial cells and incubated for 4 h at 37 °C. Cells (100 µl) were centrifuged at 3,200 g for 10 min, washed in PBS, and resuspended in 100 µl of PBS. Cells were then fixed in 4% PFA for 15 min and washed in PBS. ELMO glow discharge (Courdouan Technologies) was used to glow-discharge the carbon film for 1 min, after which 10 µl of the sample was loaded on the carbon film and excess liquid removed with filter paper. Ten microlitres of uranium acetate was added to the sample, left for 2 min, after which the samples were imaged with a JEOL 2100 transmission electron microscope.

### RNA sample preparation and sequence analysis

Overnight cultures of bacteria were diluted to 0.1 OD_600_ in caMHB, grown to mid-log phase, and diluted to 0.4 OD_600_. Equal volume (500 µl) of either Bald’s eyesalve or media (control) was added to culture at 2×MIC to make up a final volume of 1 ml. The treated cultures (*A. baumannii* ATCC 19606 and *S. aureus* Newman) were incubated for 2 h, after which the media and treatment were washed off by centrifuging at 10,000 rpm for 10 min. Cell pellets were then resuspended in PBS, centrifuged at 10,000 rpm for 10 min, snap frozen, and sent for RNA extraction and sequencing by Genewiz. Quality of the raw fastq files was checked with fastqc (68), after which trimommatic (69) was used to remove Illumina adapter sequences from the reads. Alignment was done with Bowtie2 (70). For *S. aureus*, all reads were aligned to *S. aureus* Newman whole genome sequence (accession number - NC009641.1, assembly - GCF_000010465.1), while *A. baumannii* reads were aligned to *A. baumannii* ATCC 19606 whole genome sequence (accession number - NZ_CP046654.1, assembly - GCF_009759685.1). The aligned sam files were converted to bam format using samtools (71), after which featureCounts (72) was used to obtain read counts. DESeq2 (73) in R v4.3.3 (74) was then used to determine differentially expressed genes, while Gene Ontology (GO) analysis was performed with FUNAGEpro v3 (29).

### Quorum-sensing assay

Overnight cultures of *S. aureus* RN6390 P*_xylA_*:*gfp*:*luxCDABE* (a growth-dependent reporter), *S. aureus* RN6390 P3:*gfp*:l*uxCDABE* (autoinducing peptide (AIP)-dependent reporter) (41), *P. aeruginosa* NPAO1 (Nottingham PAO1 lab strain) mini-CTX::*luxCDABE* (growth-dependent reporter), and *P. aeruginosa* NPAO1 mini-CTX::*lasB-luxCDABE* (C12-homoserine lactone-dependent reporter) (75) on LB were diluted to 0.1-0.2 OD_600_ and allowed to grow to mid-log phase in SWF, after which cells were diluted to 0.08-0.1 OD_600_. Two hundred microlitres (200 µl) of the diluted cultures were added to wells of a black 96-well flat-bottom microtitre plate CLS3596 (Corning Inc., US). After which 100 µl of treatment (sterile distilled water (control), 10% v/v Bald’s eyesalve, 1% Bald’s eyesalve, or 0.1% Bald’s eyesalve was added to the wells in triplicate. These treatments resulted in final concentrations of Bald’s eyesalve in the cultures of 3.3%, 0.33% and 0.033%, approximately corresponding to the MIC, 0.1×MIC and 0.01×MIC of *S. aureus* Newman measured in caMHB following the EUCAST protocol. The plate was incubated at 37°C for 20 h with periodic shaking in a Tecan SPARK 10 M (Tecan, Switzerland) plate reader, and the optical density (OD_600_) and luminescence of the wells (relative luminescence units, RLU) were taken every 15 minutes. Reporter gene expression was expressed as RLU/ OD_600_. None of the eyesalve treatments inhibited bacterial growth, which is expected because the assay starts with a much higher concentration of bacteria than the EUCAST MIC protocol, this inoculum is taken from an log-phase culture rather than colonies, and SWF has been shown to increase the tolerance of bacteria to Bald’s eyesalve (18).

### Biofilm inhibition assay

Triplicate cultures of *S. aureus* Newman and *A. baumannii* ATCC 19606 were grown in LB agar were grown overnight at 37°C with shaking. Cultures were resuspended in caMHB to 0.008-0.1 OD_600_ and 50 µl of each culture aliquoted into six wells of a black, transparent-bottomed 96-well plate (Corning Costar catalogue no. 3603). 50 µl of caMHB containing eyesalve at 2× MIC (6.25% v/v) was added to three wells of each culture to create a final concentration of 1× MIC (3.125% v/v). Fifty µl of caMHB was added to the other three wells of each culture to provide a no-treatment control. The optotracer EbbaBiolight 680 (Ebba Biotech AB) was added to all wells, at a final concentration of 1 µg/ml. Duplicate wells were also filled with 100 µl sterile caMHB plus 1 µg/ml EbbaBiolight 680, ± eyesalve at 1× MIC to check that no fluorescence occurred in the absence of bacteria, and single wells were filled with 100 µl sterile caMHB ± eyesalve at 1× MIC to control for any background fluorescence of the eysalve. These wells also served as sterility controls. The plate was incubated at 37°C for 48 h with periodic shaking in a Tecan SPARK 10 M (Tecan, Switzerland) plate reader, and the optical density (OD_600_) and fluorescence of the wells with ex/em 550/670 nm (relative fluorescence units, RFU) were taken every 15 minutes. Inspection of the data showed almost no change in either OD or RFU for all cultures after 24h, thus only the first 24h of data are presented.

### Long-term evolution experiment

A long-term evolution experiment was carried out to determine the resistance evolution pattern of *S. aureus*, *A. baumannii* and *P. aeruginosa* to both Bald’s eyesalve and its derived cocktail as previously described (76) with modifications. Briefly, the MICs of the treatments (Bald’s eyesalve, meropenem (for *A. baumannii* and *P. aeruginosa*), vancomycin (for *S. aureus*), and ciprofloxacin (all three)) against the bacterial strains (*S. aureus, A. baumannii,* and *P. aeruginosa*), in caMHB were first determined as previously described. Two microlitres (2 µl) of bacterial inoculum at approximately 10^5^ CFU was taken from the 0.5×MIC well and added to a new 96-well plate CL3596 (Corning Inc., US) containing 198 µl of caMHB with 0.5×MIC and 1×MIC of the treatments to make a final volume of 200 µl in each well. Three independent populations per treatment was used in this experiment. The plates were sealed with Parafilm^TM^ and incubated at 37°C for 20-24 h without shaking. For each treatment, bacteria from the well containing the highest treatment concentration with visible growth were selected for passage to fresh medium. Two microlitres (2 µl) of culture from these wells were inoculated into duplicate wells of fresh medium (198 µl) containing 1× and 2× the concentration of treatment present in the originating well. If no growth was seen in the presence of the lower concentration, the concentrations were adjusted to 0.5× the lowest concentration of the previous day. This process was repeated for 14 days. Populations were stored in 25% glycerol in LB at -80°C.

## AUTHOR CONTRIBUTIONS

Author contributions following CRedit Taxonomy:

Conceptualisation: FH, OO, JJ, CC.

Investigation: OO, JF-P, AG, BA, JR, NR, FH.

Formal analysis: OO, JF-P.

Methodology (methods development): SH, NEH, RGM.

Methodology (pilot investigation): AG, SS.

Supervision: FH, CC, SH, SPD, OO.

Writing – original draft: OO, FH.

Writing – review & editing: all authors.

## ACKNOWLEDGMENTS

We thank Christina Lee and Aled Roberts for all of the work and discussion that led to the foundation of the Ancientbiotics team and, ultimately, the present research. We thank Ian Hands-Portman and the Warwick Microscopy Bio-Analytical Shared Resource Laboratory for assistance with TEM; The Warwick Electron Microscope Research Technology Platform for use of SEM; Alan Cockayne, Kim Hardie and Kendra Rumbaugh for supplying bacterial strains; and Cerith Harries and Charlotte Curtis of the Warwick School of Life Science Media Preparation Service for supplying media and plates. This work benefited greatly from discussion and feedback on JFP and OO’s PhD theses from Antonia Sagona, Cassandra Quave, Henrik Strahl, Christopher Rodrigues and Stephen Parnell. A large number of lab members and students past and present contributed to the prior research that led to the present work, and we thank them all - especially Jason Millington and Thorulf Vargsen for contributing to a pilot resistance evolution experiment. We also gratefully acknowledge the support of the University of Warwick Institute of Advanced Studies (IAS) through an Early Career Fellowship awarded to OQO during the writing up of this manuscript.

## ETHICS APPROVAL

No ethical approval was required for this study.

