## Supplementary File for "Uncovering the mechanism of action of a historical antimicrobial against *S. aureus* and *A. baumannii*"

Oluwatosin Qawiyy Orababa *et al*.

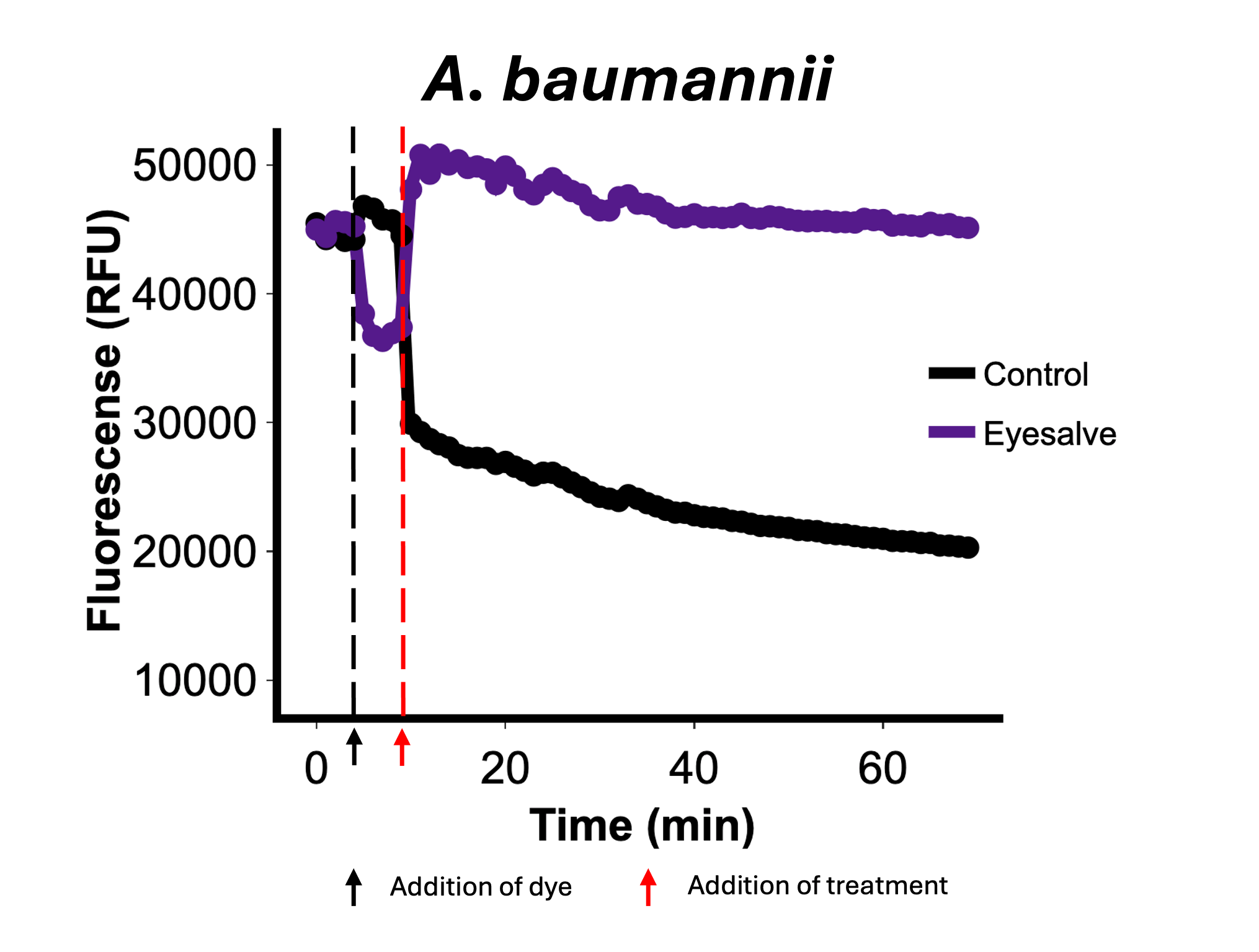

**Supplementary FIG 1**. Mid-exponential growth *A. baumannii* ATCC 19606 was diluted to 0.5 OD_600_ and pre-incubated with 30 µg/ml polymyxin B nonapeptide for 15 min and then stained with DiSC3(5) for 5 min and treated with 7.5% Bald’s Eyesalve (purple) or water (black), and their fluorescence monitored for 1 h. There was increased DiSC3(5) fluorescence, indicating membrane disruption in samples treated with Bald’s eyesalve. The red arrow indicates point of treatment.

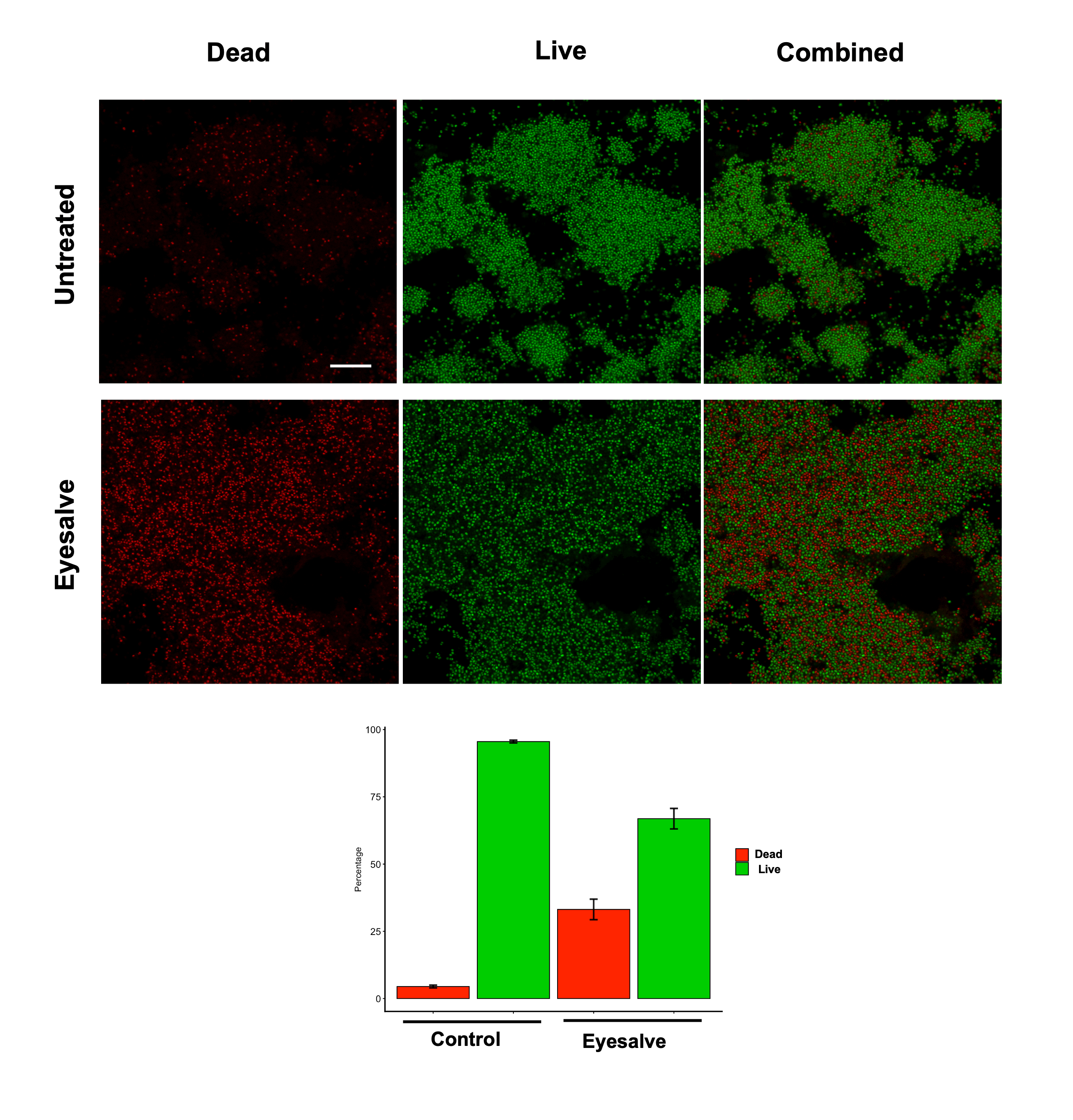

**Supplementary FIG 2.** *S. aureus* membrane permeabilisation/lysis by Bald's eyesalve. A. A 24-hour *S. aureus* wound biofilm (on the *in vitro* soft tissue model) was treated with either sterile water (control) or Bald's eyesalve overnight. This was then stained with propidium iodide and Syto-9 for 15 min in the dark and viewed under a laser scanning confocal microscope. Number of live and dead cells were counted with Biofilm viability counter on fiji and plotted. Bars are averages from 3 images. Error bars represent mean +/- standard deviation. Scale bars = 10 µm.

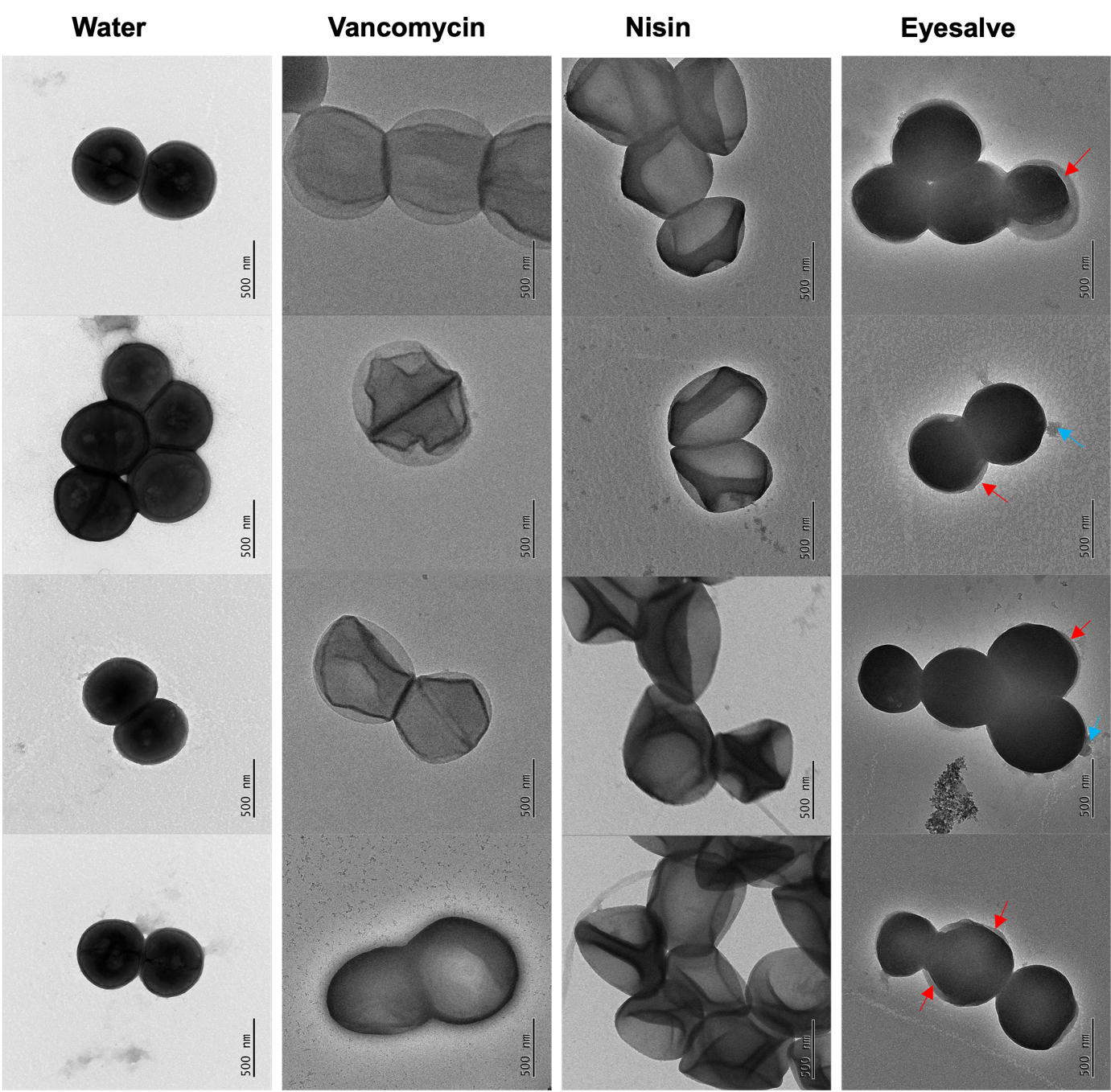

**Supplementary FIG 3. Bald’s eyesalve caused the deformation of *S. aureus* cytoplasmic membrane.** *S. aureus* Newman was treated with water, Bald’s eyesalve (100%), nisin (150 µg/ml), or vancomycin (100 µg/ml) for 4 h. Treated cells were negatively stained using ELMO glow discharge and uranium acetate. The samples were then imaged using a transmission electron microscope.

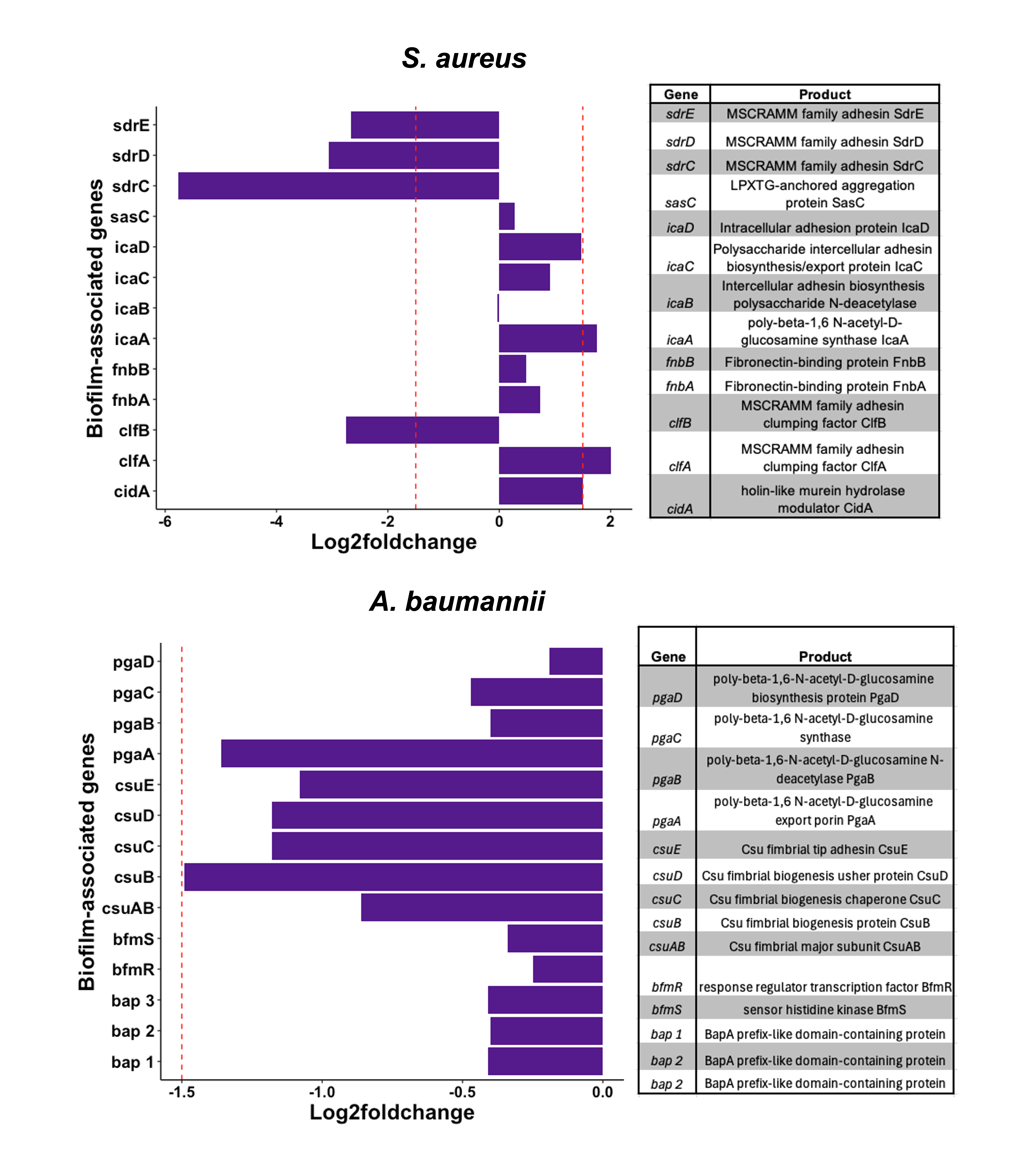

**Supplementary FIG 4. Bald’s eyesalve significantly affected the expression of some genes involved in biofilm formation in *S. aureus* and *A. baumannii*.** **A.** Expression profile of significantly differentially expressed (p < 0.05) biofilm-associated genes in *S. aureus* Newman in response to Bald’s eyesalve. Red dashed lines indicate |log2-fold change| = 1.5, n = 4 cultures per treatment. **B.** Expression profile of significantly differentially expressed (p < 0.05) biofilm-associated genes in *A. baumannii* ATCC 19606 in response to Bald’s eyesalve. Red dashed line indicates log2-fold change = -1.5, n = 3 cultures per treatment.

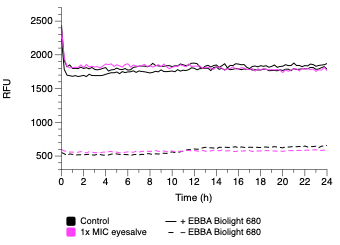

**Supplementary FIG 5.** Fluorescence of the optotracer EbbaBiolight 680 did not change over 24h of incubation at 37°C in caMHB without bacteria, and was not affected by the presence of eyesalve at 3.125% (MIC for *S. aureus* Newman and *A. baumannii* ATCC 19606).

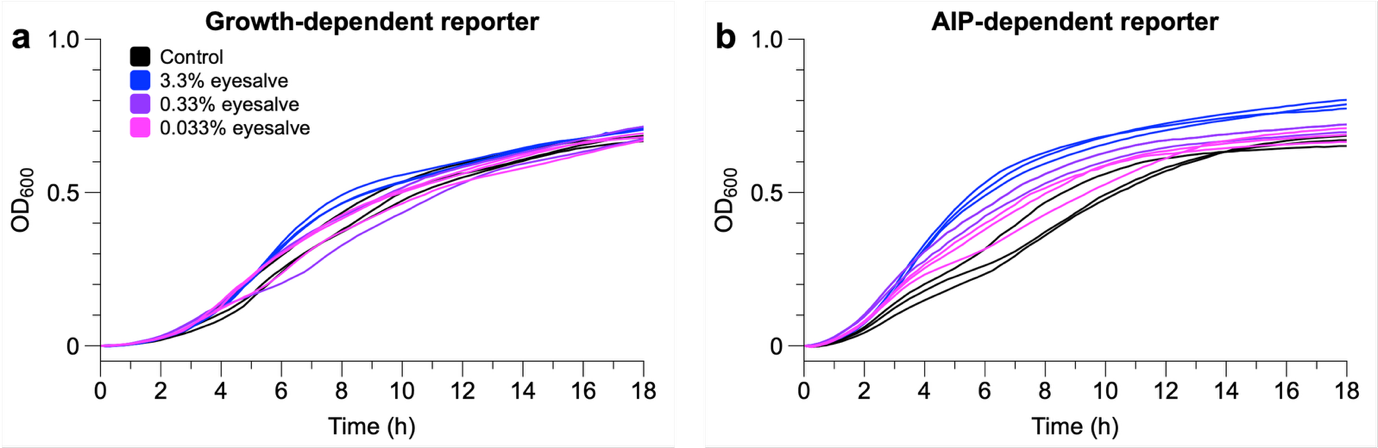

**Supplementary FIG 6.** *S. aureus* growth-dependent/constitutive reporter (**a**, P*_xylA_*:*gfp*:*luxCDABE*) and auto-inducing peptide (AIP)-dependent reporter (**b**, P3:*gfp*:l*uxCDABE*) were exposed to 3.3%, 0.33% or 0.033% v/v, or to sterile water. OD_600_ and luminescence (relative light units, RLU) were measured at intervals. Each line represents an individual bacterial culture, and OD values were standardised by subtracting the initial OD value for that population. RLU/OD data is shown in FIG 6. Exposure to eyesalve either did not noticeably affect growth or slightly enhanced it (potentially due to the presence of plant-derived carbohydrates), confirming that the concentrations used were not inhibitory under these conditions.

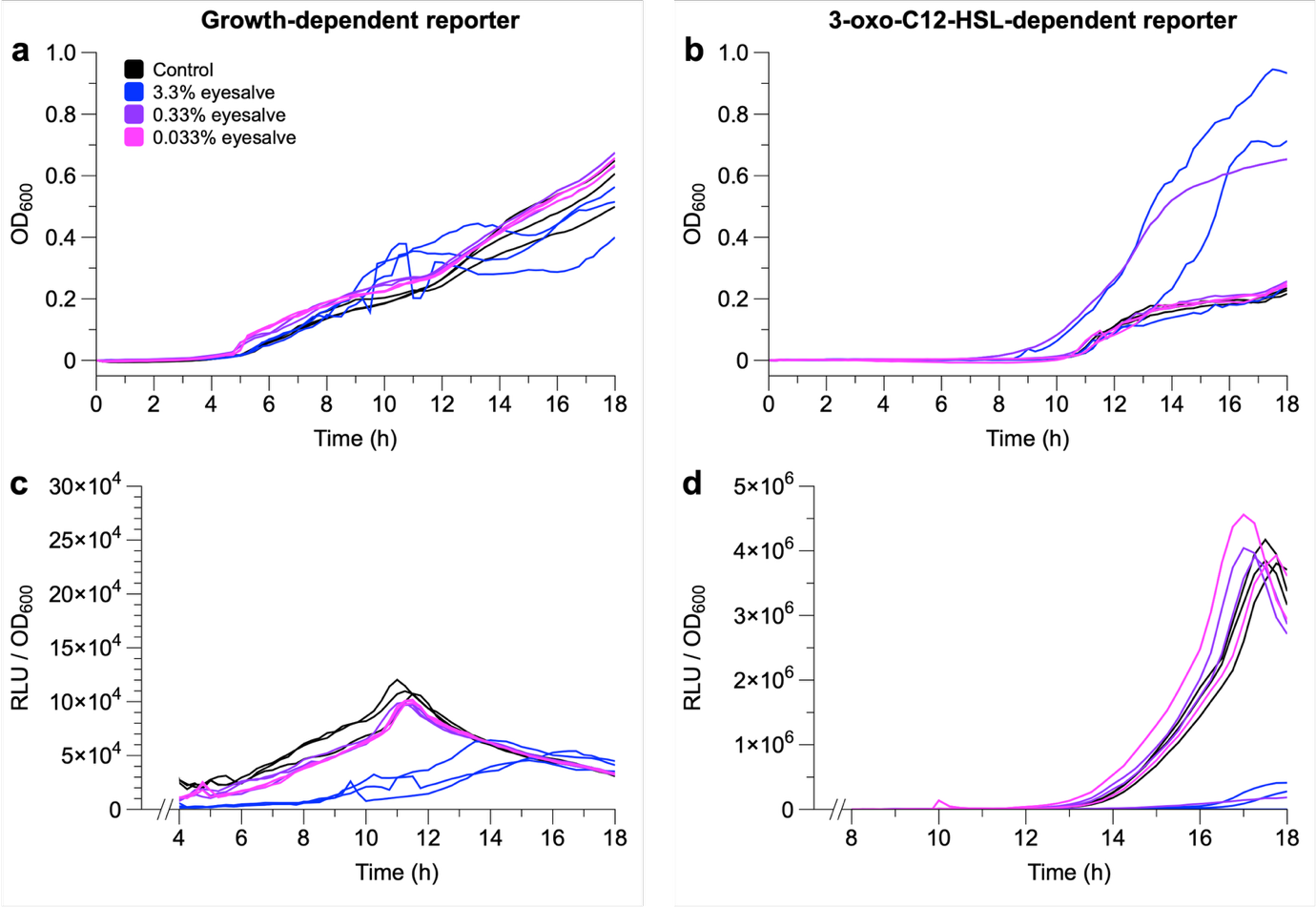

**Supplementary FIG 7.** Mid-log phase populations of *P. aeruginosa* expressing a growth-dependent/constitutive reporter (**a, c**: mini-CTX::*luxCDABE*) or a 3-oxo-C12-HSL-dependent reporter (**b, d**: mini-CTX::*lasB-luxCDABE* ) were exposed to Bald’s eyesalve concentrations of 3.3%, 0.33% or 0.033% v/v, or to sterile water. OD_600_ and luminescence (relative light units, RLU) were measured at intervals. Each line represents an individual bacterial culture, and OD values were standardised by subtracting the initial OD value for that population. **a, b)** Exposure to eyesalve either did not noticeably affect growth, or slightly enhanced it (potentially due to the presence of plant-derived carbohydrates), confirming that the concentrations used were not inhibitory under these conditions. The jaggedness and extreme OD values for some populations exposed to higher concentrations of eyesalve likely result from the turbidity of the eyesalve preparation. **c, d)** Plotting RLU/OD from the start of exponential growth phase shows that for both reporters, 3.3% eyesalve shifted the peak of luminescence right and reduced its magnitude, but lower concentrations had no discernible effect on expression. This is consistent with 3.3% eyesalve causing generalised cellular stress, and the eyesalve having no specific effect on 3-oxo-C12-HSL-mediated quorum sensing.
